# Efficient coding in biophysically realistic excitatory-inhibitory spiking networks

**DOI:** 10.1101/2024.04.24.590955

**Authors:** Veronika Koren, Simone Blanco Malerba, Tilo Schwalger, Stefano Panzeri

## Abstract

The principle of efficient coding posits that sensory cortical networks are designed to encode maximal sensory information with minimal metabolic cost. Despite the major influence of efficient coding in neuroscience, it has remained unclear whether fundamental empirical properties of neural network activity can be explained solely based on this normative principle. Here, we derive the structural, coding, and biophysical properties of excitatory-inhibitory recurrent networks of spiking neurons that emerge directly from imposing that the network minimizes an instantaneous loss function and a time-averaged performance measure enacting efficient coding. We assumed that the network encodes a number of independent stimulus features varying with a time scale equal to the membrane time constant of excitatory and inhibitory neurons. The optimal network has biologically-plausible biophysical features, including realistic integrate-and-fire spiking dynamics, spike-triggered adaptation, and a non-specific excitatory external input. The excitatory-inhibitory recurrent connectivity between neurons with similar stimulus tuning implements feature-specific competition, similar to that recently found in visual cortex. Networks with unstructured connectivity cannot reach comparable levels of coding efficiency. The optimal ratio of excitatory vs inhibitory neurons and the ratio of mean inhibitory-to-inhibitory vs excitatory-to-inhibitory connectivity are comparable to those of cortical sensory networks. The efficient network solution exhibits an instantaneous balance between excitation and inhibition. The network can perform efficient coding even when external stimuli vary over multiple time scales. Together, these results suggest that key properties of biological neural networks may be accounted for by efficient coding.

## Introduction

Information about the sensory world is represented in the brain through the dynamics of neural population activity ^1,2^. One prominent theory about the principles that may guide the design of neural computations for sensory function is efficient coding ^3,4,5^. This theory posits that neural computations are optimized to maximize the information that neural systems encode about sensory stimuli while at the same time limiting the metabolic cost of neural activity. Efficient coding has been highly influential as a normative theory of how networks are organized and designed to optimally process natural sensory stimuli in visual ^6,7,8,9,10,11^, auditory ^12^ and olfactory sensory pathways ^13^.

The first normative neural network models ^4,10^ designed with efficient coding principles had at least two major levels of abstractions. First, neural dynamics was greatly simplified, ignoring the spiking nature of neural activity. Instead, biological networks often encode information through millisecond-precise spike timing^14,15,16,17,18,19,20^. Second, these earlier contributions mostly considered encoding of static sensory stimuli, whereas the sensory environment changes continuously at multiple timescales and the dynamics of neural networks encodes these temporal variations of the environment^21,22,23,24^.

Recent years have witnessed a considerable effort and success in laying down the mathematical tools and methodology to understand how to formulate efficient coding theories of neural networks with more biological realism ^25^. This effort has established the incorporation of recurrent connectivity^26,27^, of spiking neurons, and of time-varying stimulus inputs ^28,29,30,31,32,33,34,35^. In these models, the efficient coding principle has been implemented by designing networks whose activity maximizes the encoding accuracy, by minimizing the error between a desired representation and a linear readout of network’s activity, subject to a constraint on the metabolic cost of processing. This double objective is captured by a loss function that trades off encoding accuracy and metabolic cost. The minimization of the loss function is performed through a greedy approach, by assuming that a neuron will emit a spike only if this will decrease the loss. This, in turn, yields a set of leaky integrate-and-fire (LIF) neural equations^28,29^, which can also include biologically plausible non-instantaneous synaptic delays^36,35,34^. Although most initial implementations did not respect Dale’s law, further studies analytically derived efficient networks of excitatory (E) and inhibitory (I) spiking neurons that respect Dale’s law ^28,37,31,38^ and included spike-triggered adaptation ^38^. These networks take the form of generalized leaky integrate-and-fire (gLIF) models neurons, which are realistic models of neuronal activity^39,40,41^ and capable of accurately predicting real neural spike times *in vivo* ^42^. Efficient spiking models thus have the potential to provide a normative theory of neural coding through spiking dynamics of E-I circuits ^43,38,44^ with high biological plausibility.

However, despite the major progress described above, we still lack a thorough characterization of which structural, coding, biophysical and dynamical properties of excitatory-inhibitory recurrent spiking neural networks directly relate to efficient coding. Previous studies only rarely made predictions that could be quantitatively compared against experimentally measurable biological properties. As a consequence, we still do not know which, if any, fundamental properties of cortical networks emerge directly from efficient coding.

To address the above questions, we systematically analyze our biologically plausible efficient coding model of E and I neurons that respects Dale’s law^38^. We make concrete predictions about experimentally measurable structural, coding and dynamical features of neurons that arise from efficient coding. We systematically investigate how experimentally measurable emergent dynamical properties, including firing rates, trial-to-trial spiking variability of single neurons and E-I balance ^45^, relate to network optimality. We further analyze how the organization of the connectivity arising by imposing efficient coding relates to the anatomical and effective connectivity recently reported in visual cortex, which suggests competition between excitatory neurons with similar stimulus tuning. We find that several key and robustly found empirical properties of cortical circuits match those of our efficient coding network. This lends support to the notion that efficient coding may be a design principle that has shaped the evolution of cortical circuits and that may be used to conceptually understand and interpret them.

## Results

### Assumptions and emergent structural properties of the efficient E-I network derived from first principles

We study the properties of a spiking neural network in which the dynamics and structure of the network are analytically derived starting from first principles of efficient coding of sensory stimuli. The model relies on a number of assumptions, described next.

The network responds to *M* time-varying features of a sensory stimulus, ***s***(*t*) = [*s*_1_(*t*), …, *s*_*M*_(*t*)] (e.g., for a visual stimulus, contrast, orientation, etc.) received as inputs from an earlier sensory area. We model each feature *s*_*k*_(*t*) as an independent Ornstein–Uhlenbeck (OU) processes (see Methods). The network’s objective is to compute a leaky integration of sensory features; the target representations of the network, ***x***(*t*), is defined as

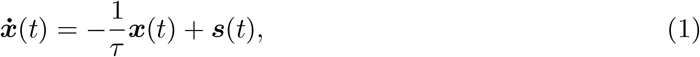

with *τ* a characteristic integration time-scale (Fig. 1A(i)). We assumed leaky integration of sensory features for consistency with previous theoretical models^37,31,33^. This assumption stems from the finding that, in many cases, integration of sensory evidence by neurons is well described by an exponential kernel ^46,47^. Additionally, a leaky integration of neural activity with an exponential kernel implemented in models of neural activity readout often explains well perceptual discrimination results^48,49,50^. This suggests that the assumption of leaky integration of sensory evidence, though possibly simplistic, captures relevant aspects of neural computations.

**Figure 1.**
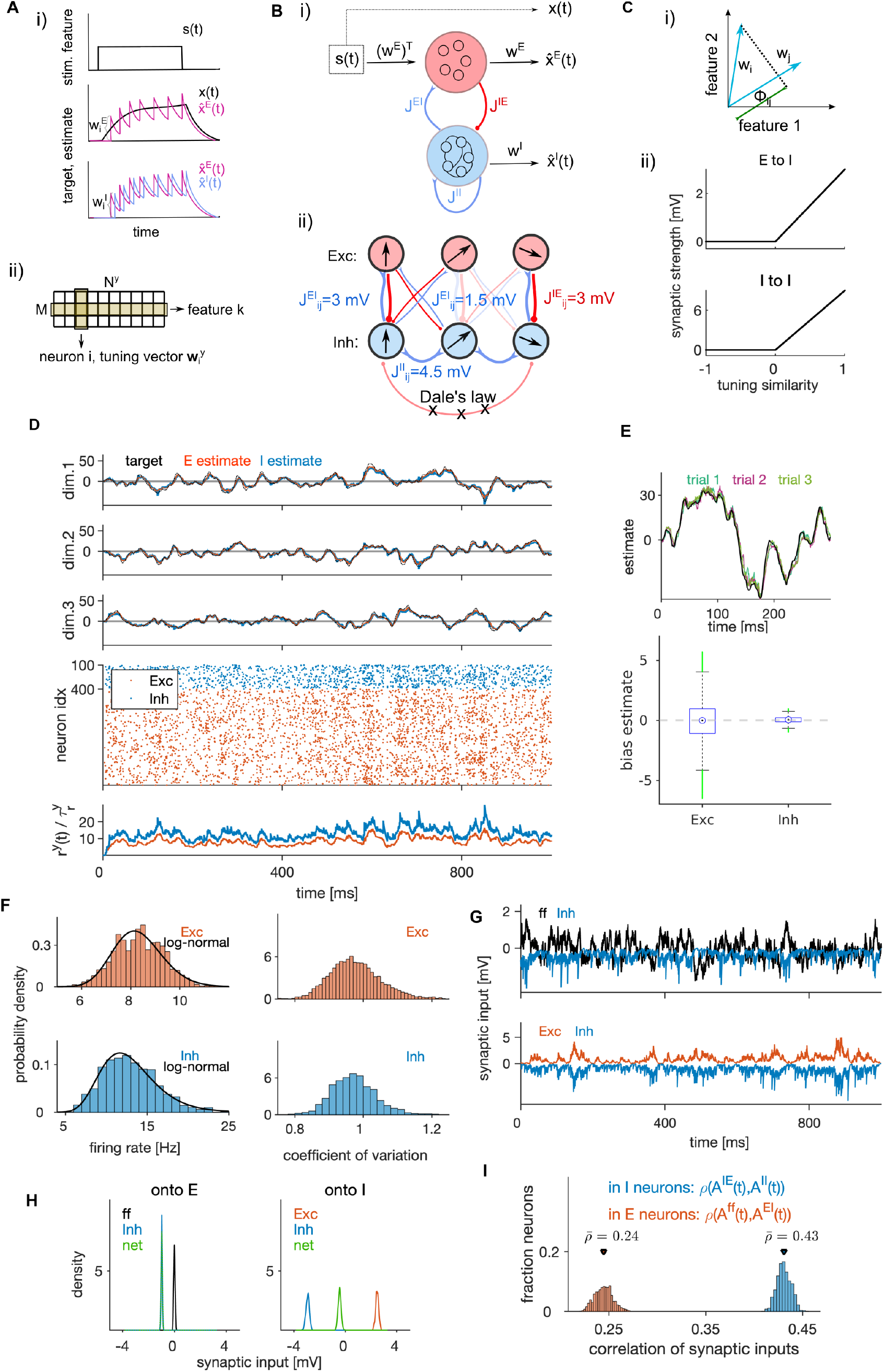
Structural and dynamical properties of the efficient E-I spiking network. **(A) (i)** Encoding of a target signal representing the evolution of a stimulus feature (top) with one E (middle) and one I spiking neuron (bottom). The target signal *x*(*t*) integrates the input signal *s*(*t*). The readout of the E neuron tracks the target signal and the readout of the I neuron tracks the readout of the E neuron. Neurons spike to bring the readout of their activity closer to their respective target. Each spike causes a jump of the readout, with the sign and the amplitude of the jump being determined by neuron’s tuning parameters. **(ii)** Schematic of the matrix of tuning parameters. Every neuron is selective to all stimulus features (columns of the matrix), and all neurons participate in encoding of every feature (rows). **(B) (i)** Schematic of the network with E (red) and I (blue) cell type. E neurons are driven by the stimulus features while I neurons are driven by the activity of E neurons. E and I neurons are connected through recurrent connectivity matrices. **(ii)** Schematic of E (red) and I (blue) synaptic interactions. Arrows represent the direction of the tuning vector of each neuron. Only neurons with similar tuning are connected and the connection strength is proportional to the tuning similarity. **(C) (i)** Schematic of similarity of tuning vectors (tuning similarity) in a 2-dimensional space of stimulus features. **(ii)** Synaptic strength as a function of tuning similarity. **(D)** Coding and dynamics in a simulation trial. Top three rows show the signal (black), the E estimate (red) and the I estimate (blue) for each of the three stimulus features. Below are the spike trains. In the bottom row, we show the average instantaneous firing rate (in Hz). **(E)** Top: Example of the target signal (black) and the E estimate in three simulation trials (colors) for one stimulus feature. Bottom: Distribution (across time) of the time-dependent bias of estimates in E and I cell type. **(F)** Left: Distribution of time-averaged firing rates in E (top) and I neurons (bottom). Black traces are fits with log-normal distribution. Right: Distribution of coefficients of variation of interspike intervals for E and I neurons. **(G)** Distribution (across neurons) of time-averaged synaptic inputs to E (left) and I neurons (right). In E neurons, the mean of distributions of inhibitory and of net synaptic inputs are very close. **(H)** Sum of synaptic inputs over time in a single E (top) and I neuron (bottom) in a simulation trial. **(I)** Distribution (across neurons) of Pearson’s correlation coefficients measuring the correlation of synaptic inputs *A*^*Ey*^ and *A*^*Iy*^ (as defined in Methods, Eq. (43)) in single E (red) and I (blue) neurons. All statistical results (E-F, H-I) were computed on 10 simulation trials of 10 second duration. For model parameters, see Table 1.

The network is composed of two neural populations of excitatory (E) and inhibitory (I) neurons, defined by their postsynaptic action which respects Dale’s law. For each population, *y* ∈ {*E, I*}, we define a population readout of each feature, 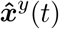, as a filtered weighted sum of spiking activity of neurons in the population,

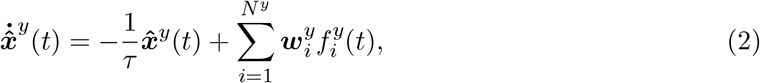

where 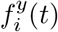 is the spike train of neuron *i* of type *y* and 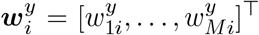 is the vector of decoding weights of the neuron for features *k* = 1, …, *M* (Fig. 1A(ii)). We assume that every neuron encodes multiple (*M >* 1) stimulus features and that the encoding of every stimulus is distributed among neurons. As a result of the optimization, the decoding weights of the neurons are equivalent to the neuron’s stimulus tuning parameters (see Methods^43^). We sampled tuning parameters uniformly from a *M*-dimensional hypersphere with unit radius, giving tuning vectors with unit length to all neurons (see Methods). To control the amount of inhibition in the network, we then multiplied the tuning vectors of I neurons with a factor *d >* 1, homogeneously across all I neurons. Normalization of decoding vectors preserves the heterogeneity of decoding weights across neurons, which may benefit coding efficiency ^51^.

Following previous work ^28,37,31^, we impose that E and I neurons have distinct normative objectives and we define specific loss functions relative to each neuron type. To implement at the same time, as requested by efficient coding, the constraints of faithful stimulus representation with limited computational resources ^52^, we define the loss functions of the population *y* ∈ {*E, I*} as a weighted sum of a time-dependent encoding error and time-dependent metabolic cost:

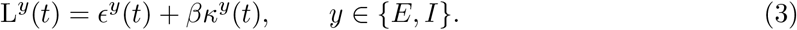

We refer to *β*, the parameter controlling the relative importance of the metabolic cost over the encoding error, as the metabolic constant of the network. We hypothesize that population readouts of E neurons, 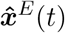, track the target representations, ***x***(*t*), and the population readouts of I neurons, 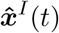, track the population readouts of E neurons, 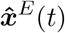, by minimizing the squared error between these quantities ^38^ (see also ^28,53^ for related approaches). Furthermore, we hypothesize the metabolic cost to be proportional to the instantaneous estimate of network’s firing frequency. We thus define the variables of loss functions in Eq. 3 as

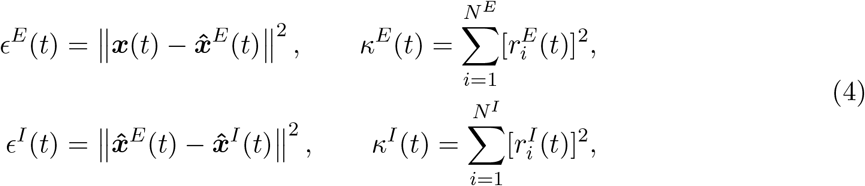

Where 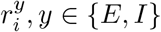, is the low-pass filtered spike train of neuron *i* (single neuron readout) with time constant 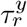, proportional to the instantaneous firing rate of the neuron: 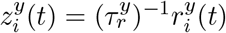. We then impose the following condition for spiking: a neuron emits a spike at time *t* only if this decreases the loss function of its population (Eq. 3) in the immediate future. The condition for spiking also includes a noise term (Methods) accounting for sources of stochasticity in spike generation ^54^ which include the effect of non-specific inputs from the rest of the brain.

We derived the dynamics and network structure of a spiking network that instantiates efficient coding (Fig. 1B, see Methods). The derived dynamics of the subthreshold membrane potential 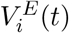 and 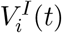 obey the equations of the generalized leaky integrate and fire (gLIF) neuron

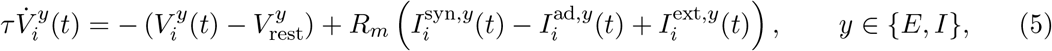

where 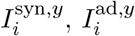, and 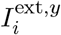 are synaptic current, spike-triggered adaptation current and nonspecific external current, respectively, *R*_*m*_ is the membrane resistance and 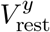 is the resting potential. This dynamics is complemented with a fire-and-reset rule: when the membrane potential reaches the firing threshold ϑ^*y*^, a spike is fired and 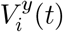 is set to the reset potential *V* ^reset,*y*^. The analytical solution in Eq. (5) holds for any number of neurons (with at least 1 neuron in each population) and predicts an optimal spike pattern to encode the presented external stimulus. Following previous work ^28^ in which physical units were assigned to derived mathematical expressions to interpret them as biophysical variables, we express computational variables (target stimuli in Eq. 1, population readouts in Eq. 2 and the metabolic constant in Eq. 3), with physical units in such a way that all terms of the biophysical model (Eq. 5) have realistic physical units.

The synaptic currents in E neurons, 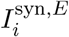, consist of feedforward currents, obtained as stimulus features ***s***(*t*) weighted by the tuning weights of the neuron, and of recurrent inhibitory currents (Fig. 1B). Synaptic currents in I neurons, 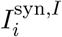, consist of recurrent excitatory and inhibitory currents. Note that there are no recurrent connections between E neurons, a consequence of our assumption of no across-feature interaction in the leaky integration of stimulus features (Eq. 8). This assumption is likely to be simplistic even for early sensory cortices ^55^. However, in other studies we found that many properties of efficient networks implementing leaky integration hold also when input features are linearly mixed during integration^38,25^.

The optimization of the loss function yielded structured recurrent connectivity (Fig. 1B(ii)-C). Synaptic strength between two neurons is proportional to their tuning similarity, forming liketo-like connectivity, if the tuning similarity is positive; otherwise the synaptic weight is set to zero (Fig. 1C (ii)) to ensure that Dale’s law is respected. A connectivity structure in which the synaptic weight is proportional to pairwise tuning similarity is consistent with some empirical observations in visual cortex ^56^ and has been suggested by previous models ^28,57^. Such connectivity organization is also suggested by across-neuron influence measured with optogenetic perturbations of visual cortex ^58,59^. While such connectivity structure is the result of optimization, the rectification of the connectivity that enforces Dale’s law does not emerge from imposing efficient coding, but from constraining the space of solutions to biologically plausible networks. Rectification also sets the overall connection probability to 0.5, which is consistent with empirically observed connection probability from pyramidal (E) neurons to parvalbumin-positive (I) neurons^60,61^, but likely overestimates the connection probability from parvalbumin-positive neurons to pyramidal neurons, which tends to be lower^61^. (For a study of how efficient coding would be implemented if the above Dale’s law constraint were removed and each neuron were free to have either an inhibitory or excitatory effect depending on the postsynaptic target, see Supplementary Text 1 and Supplementary Fig. S1A-E).

The spike-triggered adaptation current of neuron *i* in population *y*, 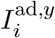, is proportional to its low-pass filtered spike train. This current realizes spike-frequency adaptation or facilitation depending on the difference between the time constants of population and single neuron readout (see Results section “Weak or no spike-triggered adaptation optimizes network efficiency”).

Finally, non-specific external currents 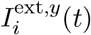 have a constant mean that depends on the parameter *β*, and fluctuations that arise from the noise with strength *σ* in the condition for spiking. The relative weight of the metabolic cost over the encoding error, *β*, controls how the network responds to feedforward stimuli, by modulating the mean of the non-specific synaptic currents incoming to all neurons. Together with the noise strength *σ*, these two parameters set the non-specific synaptic currents to single neurons that are homogeneous across the network and akin to the background synaptic input discussed in^62^. By allowing a large part of the distance between the resting potential and the threshold to be taken by the non-specific current, we found a biologically plausible set of optimally efficient model parameters (Table 1) including the firing threshold at about 20 mV from the resting potential, which is within the experimental ballpark ^63^, and average synaptic strengths of 0.75 mV (E-I and I-E synapses) and 2.25 mV (I-I synapses), which are consistent with measurements in sensory cortex ^61^. An optimal network without non-specific currents can be derived (see Methods, Eq. 25), but its parameters are not consistent with biology (see Supplementary Text 2 and Supplementary Table S1). The non-specific currents can be interpreted as synaptic currents that are modulated by larger-scale variables, such as brain states (see section “Non-specific currents regulate network coding properties”).

**Table 1.** Table of default model parameters for the efficient E-I network. Parameters above the double horizontal line are the minimal set of parameters needed to simulate model equations (Eqs. 29a-29h in Methods). Parameters below the double horizontal line are biophysical parameters, derived from the same model equations and from model parameters listed above the horizontal line. Parameters *N* ^*E*^, *M, τ* and 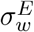 were chosen for their biological plausibility and computational simplicity. Parameters *N* ^*I*^, 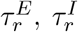, *σ*, ratio of mean E-I to I-I synaptic connectivity and *β* are parameters that maximize network efficiency (see the section “Criterion for determining model parameters” in Methods). The metabolic constant *β* and the noise strength *σ* are interpreted as global network parameters and are for this reason assumed to be the same across the E and I population, e.g., *β*^*E*^ = *β*^*I*^ = *β* and *σ*^*E*^ = *σ*^*I*^ = *σ* (see Eq. 3). The connection probability of *p*^*xy*^ = 0.5 is the consequence of rectification of the connectivity (see Eq. 24 in Methods).

| parameter | notation | value |
| --- | --- | --- |
| number of E neurons | $N^E$ | 400 |
| ratio of E to I neuron numbers | $N^E : N^I$ | 4:1 |
| number of the input features | $M$ | 3 |
| time constant of the population readout (E and I) | $\tau$ | 10 ms |
| time constant of the single neuron readout | $\tau_r^E = \tau_r^I$ | 10 ms |
| noise strength (non-specific current) | $\sigma$ | 5.0 mV |
| heterogeneity factor of tuning parameters in E | $\sigma_w^E$ | $1.0 \text{ (mV)}^{1/2}$ |
| ratio of mean I-I to E-I synaptic connectivity | mean I-I : mean E-I | 3:1 |
| metabolic constant | $\beta$ | 14 mV |
| threshold constant | $c/2$ | 18 mV |
| distance threshold to reset potential (E neurons) | $\vert \vartheta^E - V_{\text{rest}}^E\vert $ | 19 mV |
| distance threshold to reset potential (I neurons) | $\vert \vartheta^I - V_{\text{rest}}^I\vert $ | 21 mV |
| connection probability (recurrent synapses) | $p^{IE} = p^{II} = p^{EI}$ | 0.5 |
| mean E-I synaptic weight (EPSP to I at max) | $\langle J_{ij}^{IE} \rangle$ | 0.75 mV |
| mean I-E synaptic weight (IPSP to E at max) | $\langle J_{ij}^{EI} \rangle$ | 0.75 mV |
| mean I-I synaptic weight (IPSP at max) | $\langle J_{ij}^{II} \rangle$ | 2.25 mV |

To summarize, the analytical derivation of an optimally efficient network includes gLIF neurons ^64,42,41,65,66^, a distributed code with linear mixed selectivity to the input stimuli ^67,68^, spike-triggered adaptation, structured synaptic connectivity, and a non-specific external current akin to background synaptic input.

### Encoding performance and neural dynamics in an optimally efficient E-I network

The equations for the E-I network of gLIF neurons in Eq. (5) optimize the loss functions at any given time and for any set of parameters. In particular, the network equations have the same analytical form for any positive value of the metabolic constant *β*. To find a set of parameters that optimizes the overall performance, we minimized the loss function averaged over time and trials. We then optimized the parameters by setting the metabolic constant *β* such that the encoding error weights 70 % and the metabolic error weights 30 % of the average loss, and by choosing all other parameters such as to minimize numerically the average loss (see Methods). The numerical optimization was performed by simulating a model of 400 E and 100 I units, a network size relevant for computations within one layer of a cortical microcolumn ^69^. The set of model parameters that optimized network efficiency is detailed in Table 1. Unless otherwise stated, we will use the optimal parameters of Table 1 in all simulations and only vary parameters detailed in the figure axes.

With optimally efficient parameters, population readouts closely tracked the target signals (Fig. 1D, M=3, *R*^2^ = [0.95, 0.97] for E and I neurons, respectively). When stimulated by our 3-dimensional time-varying feedforward input, the optimal E-I network provided a precise estimator of target signals (Fig. 1E, top). The average estimation bias (*B*^*E*^ and *B*^*I*^, see Methods) of the network minimizing the encoding error was close to zero (*B*^*E*^ = 0.02 and *B*^*I*^ = 0.03) while the bias of the network minimizing the average loss (and optimizing efficiency) was slightly larger and negative (*B*^*E*^ = −0.15 and *B*^*I*^=−0.34), but still small compared to the stimulus amplitude (Fig. 1E, bottom, Supplementary figure S1F). Time- and trial-averaged encoding error (RMSE) and metabolic cost (MC, see Methods) were comparable in magnitude (*RMSE* = [3.5, 2.4], *MC* = [4.4, 2.8] for E and I), but with smaller error and lower cost in I, leading to a better performance in I (average loss of 2.5) compared to E neurons (average loss of 3.7). We report both the encoding error and the metabolic cost throughout the paper, so that readers can evaluate how these performance measures may generalize when weighting differently the error and the metabolic cost.

Next, we examined the emergent dynamical properties of an optimally efficient E-I network. I neurons had higher average firing rates compared to E neurons, consistently with observations in cortex ^70^. The distribution of firing rates was well described by a log-normal distribution (Fig. 1F, left), consistent with distributions of cortical firing observed empirically ^71^. Neurons fired irregularly, with mean coefficient of variation (CV) slightly smaller than 1 (Fig. 1F, right; CV= [0.97, 0.95] for E and I neurons, respectively), compatible with cortical firing ^72^. We assessed E-I balance in single neurons through two complementary measures. First, we calculated the *average* (global) balance of E-I currents by taking the time-average of the net sum of synaptic inputs (shortened to net synaptic input^73^). Second, we computed the *instantaneous* ^74^ (also termed detailed^45^) E-I balance as the Pearson correlation (*ρ*) over time of E and I currents received by each neuron (see Methods).

We observed an excess inhibition in both E and I neurons, with a negative net synaptic input in both E and I cells (Fig. 1H), indicating an inhibition-dominated network according to the criterion of average balance ^73^. In E neurons, net synaptic current is the sum of the feedforward current and recurrent inhibition and the mean of the net current is close to the mean of the inhibitory current, because feedforward inputs have vanishing mean. Furthermore, we found a moderate instantaneous balance^75^, stronger in I compared to E cell type (Fig. 1G,I, *ρ* = [0.44, 0.25], for I and E neurons, respectively), similar to levels measured empirically in rat visual cortex ^76^.

We determined optimal model parameters by optimizing one parameter at a time. To independently validate the so obtained optimal parameter set (reported in Table 1), we varied all six model parameters explored in the paper with Monte-Carlo random joint sampling (10.000 random samples), uniformly within a biologically plausible parameter range for each parameter (Table 2). We did not find any parameter configuration with lower average loss than the setting in Table 1 (Fig. 2A-B) when using the weighting of the encoding error with metabolic cost between 0.4 < *g*_*L*_ < 0.81 (Fig. 2C). The three parameter settings that came the closest to our configuration on Table 1 had stronger noise but also stronger metabolic constant than our configuration (Table 3). The second, third and fourth configurations had longer time constants of both E and I single neurons. Ratios of E-I neuron numbers and of I-I to E-I connectivity in the second, third and fourth best configuration were either jointly increased or decreased with respect to our optimal configuration. This suggests that joint covariations in parameters may influence the network’s optimality. While our finite Monte-Carlo random sampling does not fully prove the global optimality of the configuration in Table 1, it shows that it is highly efficient.

**Table 2.** Table of parameter ranges for Monte-Carlo sampling. Minimum and maximum of the uniform distributions from which we randomly drew parameters during Monte-Carlo random sampling.

| parameter | $\tau_r^E$ | $\tau_r^I$ | $\beta$ | $\sigma$ | $N^E : N^I$ | mean I-I : mean E-I |
| --- | --- | --- | --- | --- | --- | --- |
| minimum | 5 ms | 5 ms | 2 mV | 1 mV | 1 | 1 |
| maximum | 50 ms | 50 ms | 29 mV | 10 mV | 8 | 8 |

**Table 3.** Table of best four parameter settings from Monte-Carlo sampling. Best four parameter settings out of 10000 tested settings. The performance was evaluated using trial- and time-averaged loss. Each parameter setting was evaluated on 20 trials, with each trial using an independent realization of tuning parameters, noise in the non-specific current and initial conditions for the integration of the membrane potentials.

| parameter | $\tau_r^E$ | $\tau_r^I$ | $\beta$ | $\sigma$ | $N^E : N^I$ | mean I-I : mean E-I |
| --- | --- | --- | --- | --- | --- | --- |
| first | 12.6 | 11.1 | 2.1 | 4.7 | 5.4 | 3.0 |
| second | 11.4 | 10.0 | 2.9 | 6.2 | 6.1 | 3.3 |
| third | 10.0 | 10.7 | 10.1 | 3.0 | 3.2 | 2.5 |
| fourth | 12.5 | 13.5 | 2.9 | 5.4 | 4.9 | 3.5 |

**Figure 2.**
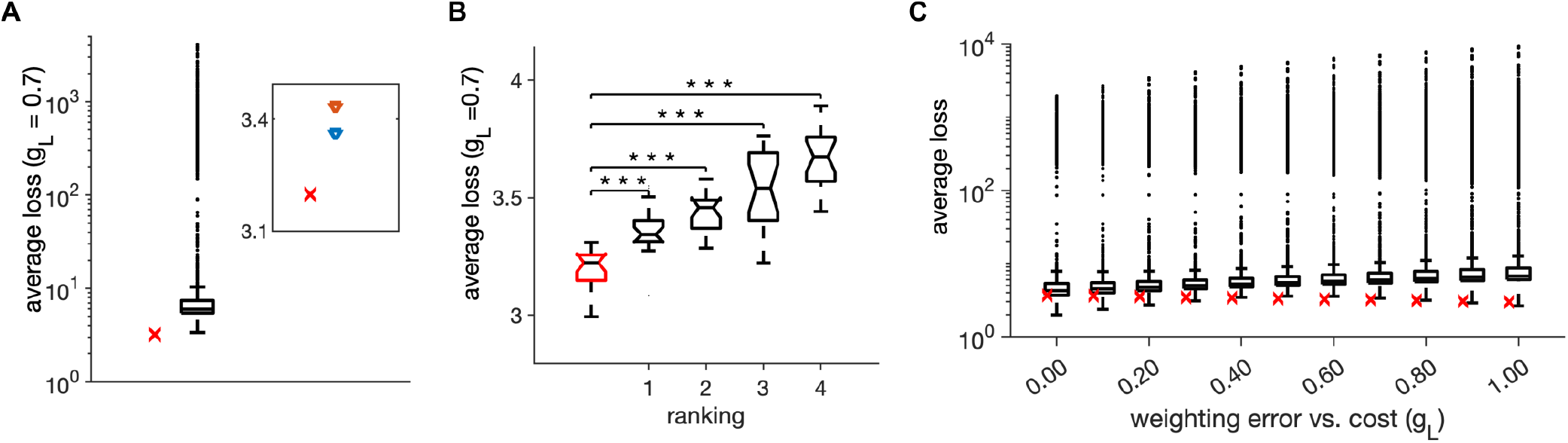
Monte-Carlo joint random sampling on six model parameters. **(A)** Distribution of the trial-averaged loss, with weighting *g*_*L*_=0.7, from 10000 random simulations and using 20 simulation trials of duration of 1 second for each parameter configuration. The red cross marks the average loss of the parameter setting in Table 1. Inset: The average loss of the parameter setting in Table 1 (red cross) and of the first- and second-best parameter settings from the random search. **(B)** Distribution of the average loss across 20 simulation trials for the parameter setting in Table 1 (red) and for the first four ranked points according to the trial-averaged loss in A. Stars indicate a significant two-tailed t-test against the distribution in red (*** indicate *p* < 0.001). **(C)** Same as in **A**, for different values of weighting of the error with the cost *g*_*L*_. Parameters for all plots are in Table 1

### Competition across neurons with similar stimulus tuning emerging in efficient spiking networks

We next explored coding properties emerging from recurrent synaptic interactions between E and I populations in the optimally efficient networks.

An approach that has recently provided empirical insight into local recurrent interactions is measuring effective connectivity with cellular resolution. Recent effective connectivity experiments photostimulated single E neurons in primary visual cortex and measured its effect on neighbouring neurons, finding that the photostimulation of an E neuron led to a decrease in firing rate of similarly tuned close-by neurons ^58^. This effective lateral inhibition ^26^ between E neurons with similar tuning to the stimulus implements competition between neurons for the representation of stimulus features ^58^. Since our model instantiates efficient coding by design and because we removed connections between neurons with different selectivity, we expected that our network implements lateral inhibition and would thus give comparable effective connectivity results in simulated photostimulation experiments.

To test this prediction, we simulated photostimulation experiments in our optimally efficient network. We first performed experiments in the absence of the feedforward input to ensure all effects are only due to the recurrent processing. We stimulated a randomly selected single target E neuron and measured the change in the instantaneous firing rate from the baseline firing rate, Δ*z*_*i*_(*t*), in all the other I and E neurons (Fig. 3A, left). The photostimulation was modeled as an application of a constant depolarising current with a strength parameter, *a*_*p*_, proportional to the distance between the resting potential and the firing threshold (*a*_*p*_ = 0 means no stimulation, while *a*_*p*_ = 1 indicates photostimulation at the firing threshold). We quantified the effect of the simulated photostimulation of a target E neuron on other E and I neurons, distinguishing neurons with either similar or different tuning with respect to the target neuron (Fig. 3A, right; Supplementary Fig. S2A-D).

**Figure 3.**
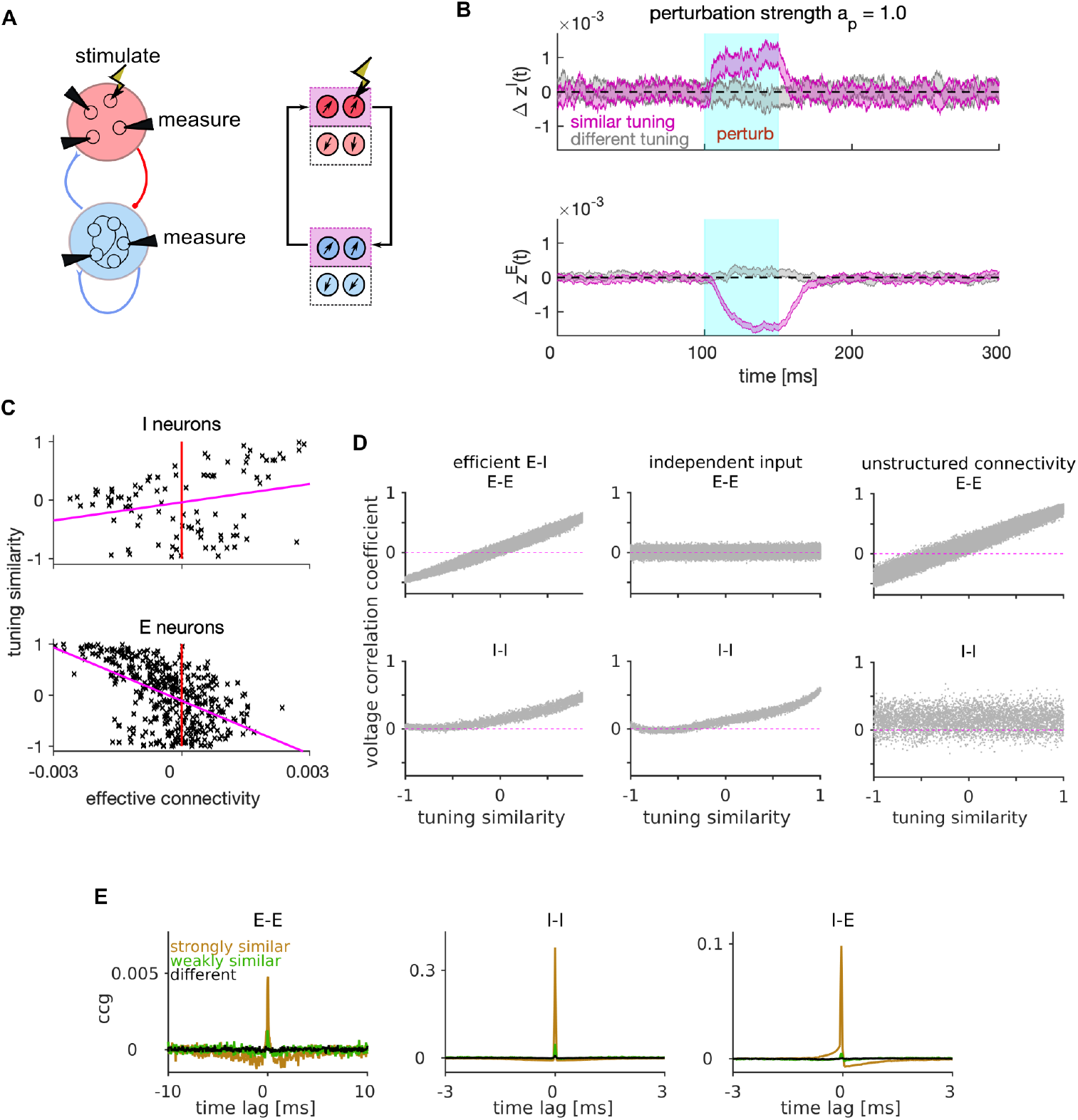
Mechanism of lateral excitation/inhibition in the efficient spiking network. **(A)** Left: Schematic of the E-I network and of the stimulation and measurement in a perturbation experiment. Right: Schematic of the propagation of the neural activity between E and I neurons with similar tuning. **(B)** Trial and neuron-averaged deviation of the firing rate from the baseline, for the population of I (top) and E (bottom) neurons with similar (magenta) and different tuning (gray) to the target neuron. The stimulation strength corresponded to an increase in the firing rate of the stimulated neuron by 28.0 Hz. **(C)** Scatter plot of the tuning similarity vs. effective connectivity to the target neuron. Red line marks zero effective connectivity and magenta line is the least-squares line. Stimulation strength was *a*_*p*_ = 1. **(D)** Correlation of membrane potentials vs. the tuning similarity in E (top) and I cell type (bottom), for the efficient E-I network (left), for the network where each E neuron receives independent instead of shared stimulus features (middle), and for the network with unstructured connectivity (right). In the model with unstructured connectivity, elements of each connectivity matrix were randomly shuffled. We quantified voltage correlation using the (zero-lag) Pearson’s correlation coefficient, denoted as 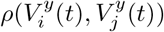, for each pair of neurons. **(E)** Average cross-correlogram (CCG) of spike timing with strongly similar (orange), weakly similar (green) and different tuning (black). Statistical results (**B-E**) were computed on 100 simulation trials. The duration of the trial in **D-E** was 1 second. Parameters for all plots are in Table 1.

The photostimulation of the target E neuron increased the instantaneous firing rate of similarly-tuned I neurons and reduced that of other similarly-tuned E neurons (Fig. 3B). We quantified the effective connectivity as the difference between the time-averaged firing rate of the recorded cell in presence or absence of the photostimulation of the targeted cell, measured during perturbation and up to 50 ms after. We found positive effective connectivity on I and negative effective connectivity on E neurons with similar tuning to the target neuron, with a positive correlation between tuning similarity and effective connectivity on I neurons and a negative correlation on E neurons (Fig. 3C). We confirmed these effects of photostimulation in presence of a weak feedforward input (Supplementary Fig. S2E), similar to the experiments of Ref^58^ in which photostimulation was applied during the presentation of visual stimuli with weak contrast. Thus, the optimal network replicates the preponderance of negative effective connectivity between E neurons and the dependence of its strength on tuning similarity found in ^58^.

In summary, lateral excitation of I neurons and lateral inhibition of E neurons with similar tuning is an emerging coding property of the efficient E-I network, which recapitulates competition between neurons with similar stimulus tuning found in visual cortex ^58,59^. An intuition of why this competition implements efficient coding is that the E neuron that fires first activates I neurons with similar tuning. In turn, these I neurons inhibit all similarly tuned E neurons (Fig. 3A, right), preventing them to generate redundant spikes to encode the sensory information that has already been encoded by the first spike. Suppression of redundant spiking reduces metabolic cost without reducing encoded information ^28,36^.

While perturbing the activity of E neurons in our model qualitatively reproduces empirically observed lateral inhibition among E neurons ^58,59^, these experiments have also reported positive effective connectivity between E neurons with very similar stimulus tuning. Our intuition is that our simple model cannot reproduce this finding because it lacks E-E connectivity.

To explore further the consequences of E-I interactions for stimulus encoding, we next investigated the dynamics of lateral inhibition in the optimal network driven by the feedforward sensory input but without perturbing the neural activity. Previous work has established that efficient spiking neurons may present strong correlations in the membrane potentials, but only weak correlations in the spiking output, because redundant spikes are prevented by lateral inhibition ^28,32^. We investigated voltage correlations in pairs of neurons within our network as a function of their tuning similarity. Because the feedforward inputs are shared across E neurons and weighted by their tuning parameters, they cause strong positive voltage correlations between E-E neuronal pairs with very similar tuning and strong negative correlations between pairs with very different (opposite) tuning (Fig. 3D, top-left). Voltage correlations between E-E pairs vanished regardless of tuning similarity when we made the feedforward inputs independent across neurons (Fig. 3D, top-middle), showing that the dependence of voltage correlations on tuning similarity occurs because of shared feedforward inputs. In contrast to E neurons, I neurons do not receive feedforward inputs and are driven only by similarly tuned E neurons (Fig. 3A, right). This causes positive voltage correlations in I-I neuronal pairs with similar tuning and vanishing correlations in neurons with different tuning (Fig. 3D, bottom-left). Such dependence of voltage correlations on tuning similarity disappears when removing the structure from the E-I synaptic connectivity (Fig. 3D, bottom-right).

In contrast to voltage correlations, and as expected by previous studies ^28,32^, the coordination of spike timing of pairs of E neurons (measured with cross-correlograms or CCGs) was very weak (Fig. 3E). For I-I and E-I neuronal pairs, the peaks of CCGs were stronger than those observed in E-E pairs, but they were present only at very short lags (lags < 1 ms). This confirms that recurrent interactions of the efficient E-I network wipe away the effect of membrane potential correlations at the spiking output level, and shows information processing with millisecond precision in these networks ^28,32,36^.

### The effect of structured connectivity on coding efficiency and neural dynamics

The analytical solution of the optimally efficient E-I network predicts that recurrent synaptic weights are proportional to the tuning similarity between neurons. We next investigated the role of such connectivity structure by comparing the behavior of an efficient network with an unstructured E-I network, similar to the type studied in previous works ^77,78,23^. We removed the connectivity structure by randomly permuting synaptic weights across neuronal pairs (see Methods). Such shuffling destroys the relationship between tuning similarity and synaptic strength (as shown in Fig. 1C(ii)) while it preserves Dale’s law and the overall distribution of connectivity weights.

We found that shuffling the connectivity structure significantly altered the efficiency of the network (Fig. 4A-B), neural dynamics (Fig. 4C-D, F-H) and lateral inhibition (Fig. 4I). In particular, structured networks differ from unstructured ones by showing better encoding performance (Fig. 4A), lower metabolic cost (Fig. 4B), weaker variance of the membrane potential over time (Fig. 4C), lower firing rates (Fig. 4D) and weaker average (Fig. 4F) and instantaneous balance (Fig. 4G) of synaptic inputs. However, we found only a small difference in the variability of spiking between structured and unstructured networks (Fig. 4E). While these results are difficult to test experimentally due to the difficulty of manipulating synaptic connectivity structures *in vivo*, they highlight the importance of the connectivity structure for cortical computations.

**Figure 4.**
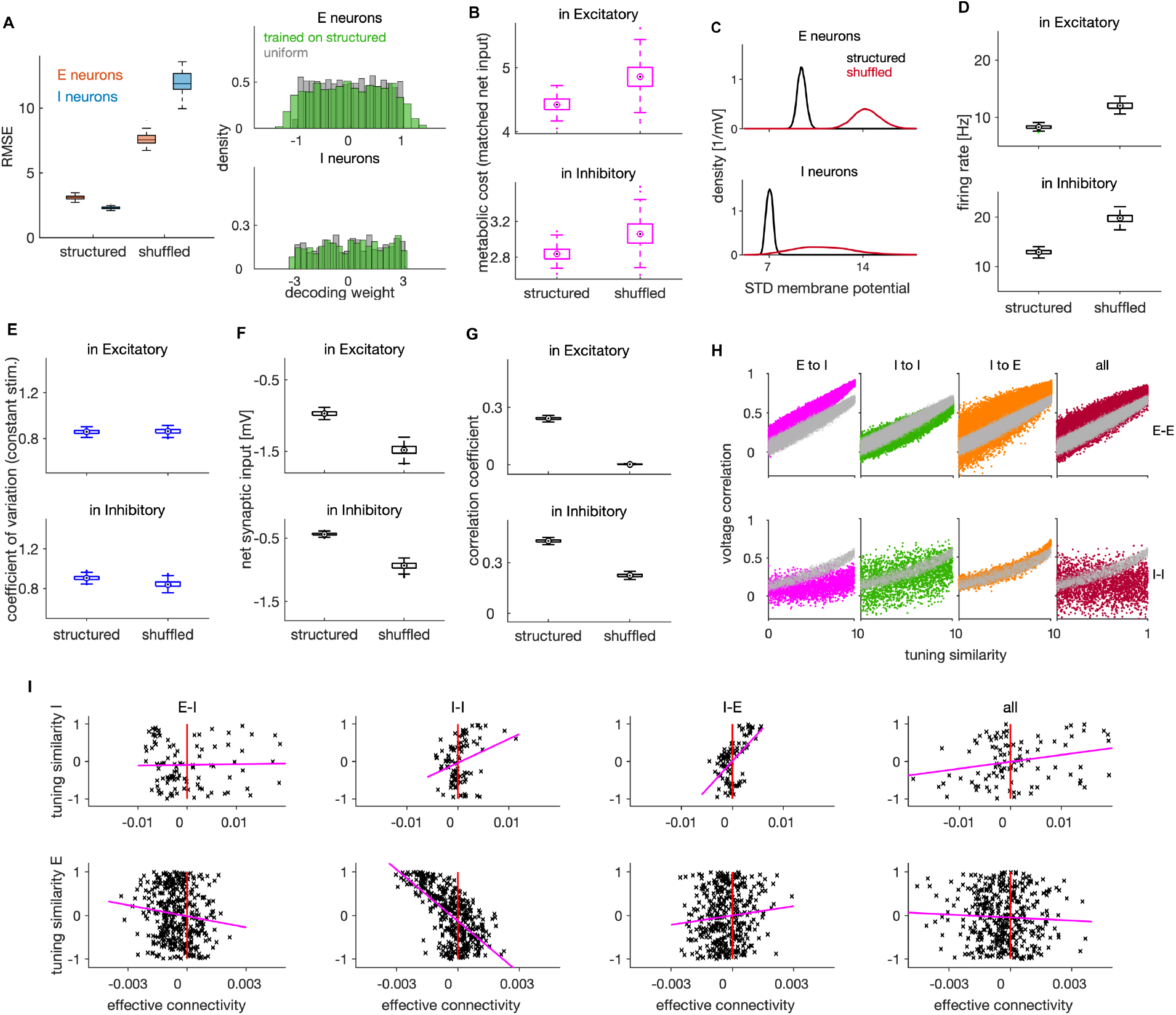
Effects of connectivity structure on coding efficiency, neural dynamics and lateral inhibition. **(A)** Left: Root mean squared error (RMSE) in networks with structured and randomly shuffled recurrent connectivity. Random shuffling consisted of a random permutation of the elements within each of the three (E-I, I-I, I-E) connectivity matrices. Right: Distribution of decoding weights after training the decoder on neural activity from the structured network (green), and a sample from uniform distribution as typically used in the optimal network. **(B)** Metabolic cost in structured and shuffled networks with matched average balance. The average balance of the shuffled network was matched with the one of the structured network by changing the following parameters: 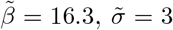 and by decreasing the amplitude of the OU stimulus by factor of 0.88. **(C)** Standard deviation of the membrane potential (in mV) for networks with structured and unstructured connectivity. Distributions are across neurons. **(D)** Average firing rate of E (top) and I neurons (bottom) in networks with structured and unstructured connectivity. **(E)** Same as in **D**, showing the coefficient of variation of spiking activity in a network responding to a constant stimulus. **(F)** Same as in **D**, showing the average net synaptic input, a measure of average imbalance. **(G)** Same as in **D**, showing the time-dependent correlation of synaptic inputs, a measure of instantaneous balance. **(H)** Voltage correlation in E-E (top) and I-I neuronal pairs (bottom) for the four cases of unstructured connectivity (colored dots) and the equivalent result in the structured network (grey dots). We show the results for pairs with similar tuning. **(I)** Scatter plot of effective connectivity versus tuning similarity to the photostimulated E neuron in shuffled networks. The title of each plot indicates the connectivity matrix that has been shuffled. The magenta line is the least-squares regression line and the photostimulation is at threshold (*a*_*p*_ = 1.0). Results were computed using 200 (**A-G**) and 100 (**H-I**) simulation trials of 1 second duration. Parameters for all plots are in Table 1.

We also compared structured and unstructured networks about their relation between pairwise voltage correlations and tuning similarity, by randomizing connections within a single connectivity type (E-I, I-I or I-E) or within all these three connectivity types at once (“all”). We found the structure of E-I connectivity to be crucial for the linear relation between voltage correlations and tuning similarity in pairs of I neurons (Fig. 4H, magenta).

Finally, we analyzed how the structure in recurrent connectivity influences lateral inhibition that we observed in efficient networks. We found that the dependence of lateral inhibition on tuning similarity vanishes when the connectivity structure is fully removed (Fig. 4I, “all” on the right plot), thus showing that connectivity structure is necessary for lateral inhibition. While networks with unstructured E-I and I-E connectivity still show inhibition in E neurons upon single neuron photostimulation (because of the net inhibitory effect of recurrent connectivity; Supplementary Fig. S3F), this inhibition was largely unspecific to tuning similarity (Fig. 4I, “E-I” and “I-E”). Unstructured connectivity decreased the correlation between tuning similarity and effective connectivity from *r* = [0.31, −0.54] in E and I neurons in a structured network to *r* = [0.02, −0.13] and *r* = [0.57, 0.11] in networks with unstructured E-I and I-E connectivity, respectively. Removing the structure in I-I connectivity, in contrast, increased the correlation between effective connectivity and tuning similarity in E neurons (*r* = [0.30, −0.65], Fig. 4I, second from the left), showing that lateral inhibition takes place irrespectively of the I-I connectivity structure.

Previous empirical ^56^ and theoretical work has established the necessity of strong E-I-E synaptic connectivity for lateral inhibition ^57,79^. To refine this understanding, we asked what is the minimal connectivity structure necessary to qualitatively replicate empirically observed lateral inhibition. We did so by considering a simpler connectivity rule than the one obtained from first principles. We assumed neurons to be connected (with random synaptic efficacy) if their tuning vectors are similar 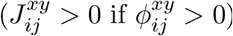 and unconnected otherwise 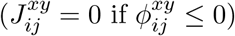, relaxing the precise proportionality relationship between tuning similarity and synaptic weights (as on Fig. 1C(ii)). We found that networks with such simpler connectivity respond to activity perturbation in a qualitatively similar way as the optimal network (Supplementary Fig. S2F) and still replicate experimentally observed activity profiles in ^58^.

While optimally structured connectivity predicted by efficient coding is biologically plausible, it may be difficult to realise it exactly on a synapse-by-synapse basis in biological networks. Following ^80^, we verified the robustness of the model to small deviations from the optimal synaptic weights by adding a random jitter, proportional to the synaptic strength, to all synaptic connections (see Methods). The encoding performance and neural activity were barely affected by weak and moderate levels of such perturbation (Supplementary Fig. S3 G-H), demonstrating that the network is robust against random jittering of the optimal synaptic weights.

In summary, we found that some aspects of recurrent connectivity structure, such as the liketo-like organization, are crucial to achieve efficient coding. Instead, for other aspects there is considerable flexibility; the proportionality between tuning similarity and synaptic weights is not crucial for efficiency and small random jitter of optimal weights has only minor effects. Structured E-I and I-E, but not I-I connectivity, is necessary for implementing experimentally observed pattern of lateral inhibition whose strength is modulated by tuning similarity.

### Weak or no spike-triggered adaptation optimizes network efficiency

We next investigated the role of within-neuron feedback triggered by each spike, 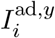, that emerges from the optimally efficient solution (Eq. 5). A previous study ^33^ showed that spike-triggered adaptation, together with structured connectivity, redistributes the activity from highly excitable neurons to less excitable neurons, leaving the population readout invariant. Here, we address model efficiency in presence of adapting or facilitating feedback as well as differential effects of adaptation in E and I neurons.

The spike-triggered within-neuron feedback 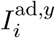 has a time constant equal to that of the single neuron readout 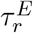 (E neurons) and 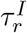 (I neurons). The strength of the current is proportional to the difference in inverse time constants of single neuron and population readouts, 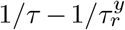. This spike-triggered current is negative, giving spike-triggered adaptation ^40^, if the single-neuron readout has longer time constant than the population readout 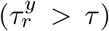, or positive, giving spike-triggered facilitation, if the opposite is true 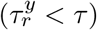 (Table 4). We expected that network efficiency would benefit from spike-triggered adaptation, because accurate encoding requires fast temporal dynamics of the population readouts, to capture fast fluctuations in the target signal, while we expect a slower dynamics in the readout of single neuron’s firing frequency, 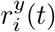, a process that could be related to homeostatic regulation of single neuron’s firing rate^81,82^. In our optimal E-I network we indeed found that optimal coding efficiency is achieved in absence of within-neuron feedback or with weak adaptation in both cell types (Fig. 5A). The optimal set of time constants 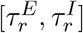 only weakly depended on the weighting of the encoding error with the metabolic cost *g*_*L*_ (Supplementary Fig. S4A). We note that adaptation in E neurons promotes efficient coding because it enforces every spike to be error-correcting, while a spike-triggered facilitation in E neurons would lead to additional spikes that might be redundant and reduce network efficiency. Contrary to previously proposed models of adaptation in LIF neurons ^39,83^, the strength and the time constant of adaptation in our model are not independent, but they both depend on 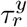, with larger 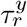 yielding both longer and stronger adaptation.

**Table 4.** Relation of time constants of single-neuron and population readout set an adaptation or a facilitation current. The population readout that evolves on a faster (slower) time scale than the single neuron readout determines a spike-triggered adaptation (facilitation) in its own cell type.

| relative speed | relation of time constants | current |
| --- | --- | --- |
| $\hat{x}^E$ faster than $r^E$ | $\tau < \tau_r^E$ | adaptation in E |
| $\hat{x}^E$ slower than $r^E$ | $\tau > \tau_r^E$ | facilitation in E |
| $\hat{x}^I$ faster than $r^I$ | $\tau < \tau_r^I$ | adaptation in I |
| $\hat{x}^I$ slower than $r^I$ | $\tau > \tau_r^I$ | facilitation in I |

**Figure 5.**
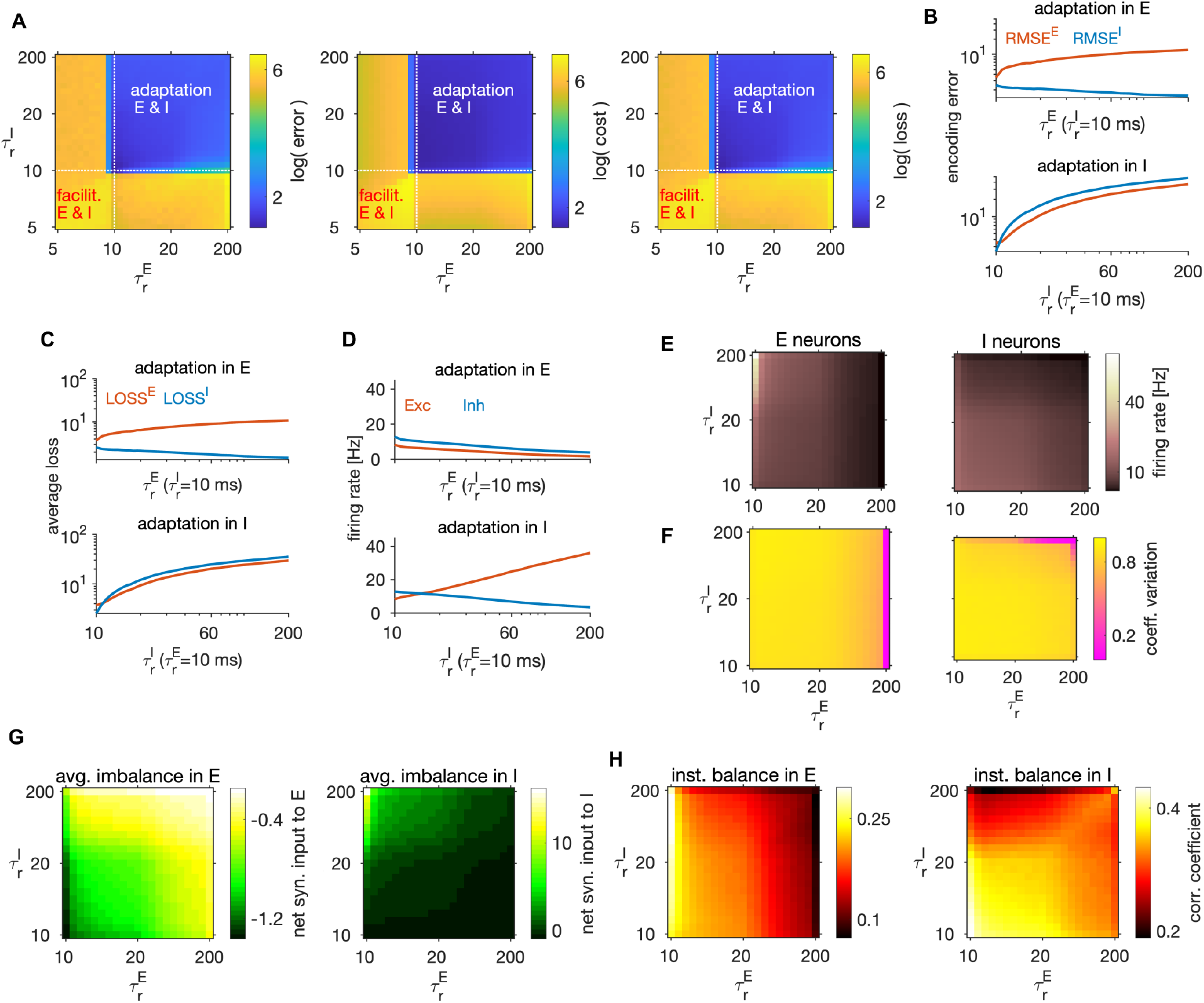
Adaptation, network coding efficiency and excitation-inhibition balance. **(A)** The encoding error (left), metabolic cost (middle) and average loss (right) as a function of single neuron time constants 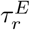 and 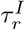, in units of ms. These parameters set the sign, the strength, as well as the time constant of the feedback current in E and I neurons. Best performance (lowest average loss) is obtained in the top right quadrant, where the feedback current is spike-triggered adaptation in both E and I neurons. The performance measures are computed as a weighted sum of the respective measures across the E and I populations with equal weighting for E and I. All measures are plotted on the scale of the natural logarithm for better visibility. **(B)** Top: Log-log plot of the RMSE of the E (red) and the I (blue) estimates as a function of the time constant of the single neuron readout of E neurons, 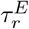, in the regime with spike-triggered adaptation. Feedback current in I neurons is set to 0. Bottom: Same as on top, as a function of 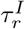 while the feedback current in E neurons is set to 0. **(C)** Same as in **B**, showing the average loss. **(D)** Same as in **B**, showing the firing rate. **(E)** Firing rate in E (left) and I neurons (right), as a function of time constants 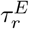 and 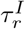. **(F)** Same as in **E**, showing the coefficient of variation. **(G)** Same as **E**, showing the average net synaptic input, a measure of average imbalance. **(H)** Same as **E**, showing the average net synaptic input, a measure of instantaneous balance. All statistical results were computed on 100 simulation trials of 1 second duration. For other parameters, see Table 1.

To gain insights on the differential effect of adaptation in E vs I neurons, we set the adaptation in one cell type to 0 and vary the strength of adaptation in the other cell type by varying the time constant of the single neuron readout. With adaptation in E neurons (and no adaptation in I), we observed a slow increase of the encoding error in E neurons, while the encoding error increased faster with adaptation in I neurons (Fig. 5B). Similarly, network efficiency increased slowly with adaptation in E and faster with adaptation in I neurons (Fig. 5C), thus showing that adaptation in E neurons decreases less the performance compared to the adaptation in I neurons. With increasing adaptation in E neurons, the firing rate in E neurons decreased (Fig. 5D), leading to E estimates with smaller amplitude. Because E estimates are target signals for I neurons and because weaker E signals imply weaker drive to I neurons, average loss of the I population decreased by increasing adaptation in E neurons (Fig. 5C top, blue trace).

Firing rates and variability of spiking were sensitive to the strength of adaptation. As expected, adaptation in E neurons caused a decrease in the firing levels in both cell types (Fig. 5D-E). In contrast, adaptation in I neurons decreased the firing rate in I neurons, but increased the firing rate in E neurons, due to a decrease in the level of inhibition. Furthermore, adaptation decreased the variability of spiking, in particular in the cell type with strong adaptation (Fig. 5F), a wellknown effect of spike-triggered adaptation in single neurons ^83^.

In regimes with adaptation, time constants of single neuron readout 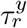 influenced the average balance (Fig. 5G) as well as the instantaneous balance (Fig. 5H) in E and I cell type. To gain a better understanding of the relationship between adaptation, E-I interactions and network optimality, we measured the instantaneous and time-averaged E-I balance while varying the adaptation parameters and studied their relation with the loss. By increasing adaptation in E neurons, the average imbalance got weaker in E neurons (Fig. 5G, left), but stronger in I neurons (Fig. 5G, right). Regimes with precise average balance in both cell types were suboptimal (compare Fig. 5A, right and G), while regimes with precise instantaneous balance were highly efficient (compare Fig. 5A, right and H).

To test how well the average balance and the instantaneous balance of synaptic inputs predict network efficiency, we concatenated the column-vectors of the measured average loss and of the average imbalance in each cell type and computed the Pearson correlation between these quantities. The correlation between the average imbalance and the average loss was weak in the E cell type (*r* = 0.16) and close to zero in the I cell type (*r* = 0.02), suggesting almost no relation between efficiency and average imbalance. In contrast, the average loss was negatively correlated with the instantaneous balance in both E (*r* = −0.35) and in I cell type (*r* = −0.45), showing that instantaneous balance of synaptic inputs is positively correlated with network efficiency. When measured for varying levels of spike-triggered adaptation, unlike the average balance of synaptic inputs, the instantaneous balance is thus mildly predictive of network efficiency.

In sum, our results show that the optimally efficient solution does not include within-neuron feedback, while a model with weak and short-lasting spike-triggered adaptation is slightly suboptimal, though still highly efficient. Our results predict that information coding would be more efficient with adaptation than with facilitation. Assuming that our I neurons describe parvalbumin-positive interneurons, our results suggest that the weaker adaptation in I compared to E neurons, reported empirically ^60^, may be beneficial for the network’s encoding efficiency.

Spike-triggered adaptation in our model captures adaptive processes in single neurons that occur on time scales lasting from a couple of milliseconds to tens of milliseconds after each spike. However, spiking in biological neurons triggers adaptation on multiple time scales, including much slower time scales on the order of seconds or tens of seconds ^84^. Our model does not capture adaptive processes on these longer time scales (but see ^33^).

### Non-specific currents regulate network coding properties

In our derivation of the optimal network, we obtained a non-specific external current (in the following, non-specific current) 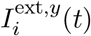. Non-specific current captures all synaptic currents that are unrelated and unspecific to the stimulus features. This non-specific term collates effects of synaptic currents from neurons untuned to the stimulus^85,86^, as well as synaptic currents from other brain areas. It can be conceptualized as the background synaptic activity that provides a large fraction of all synaptic inputs to both E and I neurons in cortical networks ^87^, and which may modulate feedforward-driven responses by controlling the distance between the membrane potential and the firing threshold^62^. Likewise, in our model, the non-specific current does not directly convey information about the feedforward input features, but influences the network dynamics.

Non-specific current comprises mean and fluctuations (see Methods). The mean is proportional to the metabolic constant *β* and its fluctuations reflect the noise that we included in the condition for spiking. Since *β* governs the trade-off between encoding error and metabolic cost (Eq. 3), higher values of *β* imply that more importance is assigned to the metabolic efficiency than to coding accuracy, yielding a reduction in firing rates. In the expression for the non-specific current, we found that the mean of the current is negatively proportional to the metabolic constant *β* (see Methods). Because the non-specific current is typically depolarizing, this means that increasing *β* yields a weaker non-specific current and increases the distance between the mean membrane potential and the firing threshold. Thus, an increase of the metabolic constant is expected to make the network less responsive to the feedforward signal.

We found the metabolic constant *β* to significantly influence the spiking dynamics (Fig. 6A). The optimal efficiency was achieved for non-zero levels of the metabolic constant (Fig. 6B), with the mean of the non-specific current spanning more than half of the distance between the resting potential and the threshold (Table 1). Stronger weighting of the loss of I compared to E neurons and stronger weighting of the error compared to the cost yielded weaker optimal metabolic constant (Supplementary Fig. S4B). Metabolic constant modulated the firing rate as expected, with the firing rate in E and I neurons decreasing with the increasing of the metabolic constant (Fig. 6C, top). It also modulated the variability of spiking, as increasing the metabolic constant decreased the variability of spiking in both cell types (Fig. 6C, bottom). Furthermore, it modulated the average balance and the instantaneous balance in opposite ways: larger values of *β* led to regimes that had stronger average balance, but weaker instantaneous balance (Fig. 6D). We note that, even with suboptimal values of the metabolic constant, the neural dynamics remained within biologically relevant ranges.

**Figure 6.**
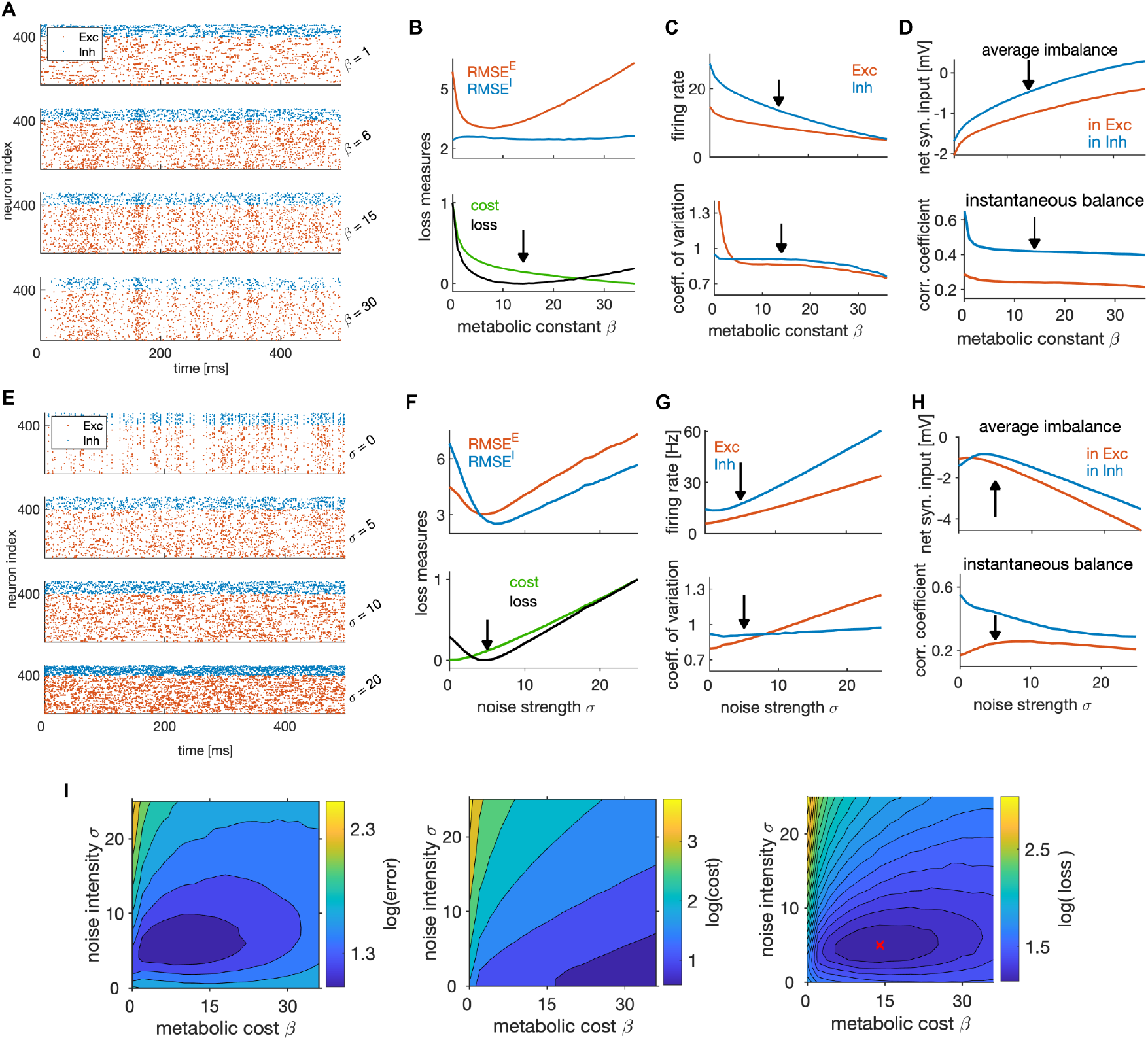
State-dependent coding and dynamics are controlled by non-specific currents. **(A)** Spike trains of the efficient E-I network in one simulation trial, with different values of the metabolic constant *β*. The network received identical stimulus across trials. **(B)** Top: RMSE of E (red) and I (blue) estimates as a function of the metabolic constant. Bottom: Normalized average metabolic cost and average loss as a function of the metabolic constant. Black arrow indicates the minimum loss and therefore the optimal metabolic constant. **(C)** Average firing rate (top) and the coefficient of variation of the spiking activity (bottom), as a function of the metabolic constant. Black arrow marks the metabolic constant leading to optimal network efficiency in **B.(D)** Average imbalance (top) and instantaneous balance (bottom) balance as a function of the metabolic constant. **(E)** Same as in **A**, for different values of the noise strength *σ*. **(F)** Same as in **B**, as a function of the noise strength. The noise is a Gaussian random process, independent over time and across neurons. **(G)** Same as **C**, as a function of the noise strength. **(H)** Top: Same as in **D**, as a function of the noise strength. **(I)** The encoding error measured as RMSE (left), the metabolic cost (middle) and the average loss (right) as a function of the metabolic constant *β* and the noise strength *σ*. Metabolic constant and noise strength that are optimal for the single parameter search (in **B** and **F**) are marked with a red cross in the figure on the right. For plots in **B-D** and **F-I**, we computed and averaged results over 100 simulation trials with 1 second duration. For other parameters, see Table 1.

The fluctuation part of the non-specific current, modulated by the noise strength *σ* that we added in the definition of spiking rule for biological plausibility (see Methods), strongly affected the neural dynamics as well (Fig. 6E). The optimal performance was achieved with non-vanishing noise levels (Fig. 6F), similarly to previous work showing that the noise prevents excessive network synchronization that would harm performance ^31,36,88^. The optimal noise strength depended on the weighting of the error with the cost, with strong weighting of the error predicting stronger noise (Supplementary Fig. S4C).

The average firing rate of both cell types, as well as the variability of spiking in E neurons, increased with noise strength (Fig. 6G), and some level of noise in the non-specific inputs was necessary to establish the optimal level of spiking variability. Nevertheless, we measured significant levels of spiking variability already in the absence of noise, with a coefficient of variation of about 0.8 in E and 0.9 in I neurons (Fig. 6G, bottom). This indicates that the recurrent network dynamics generates substantial variability even in absence of an external source of noise. The average and instantaneous balance of synaptic currents exhibited a non-linear behavior as a function of noise strength (Fig. 6H). Due to decorrelation of membrane potentials by the noise, instantaneous balance in I neurons decreased with increasing noise strength (Fig. 6H, bottom).

Next, we investigated the joint impact of the metabolic constant and the noise strength on network optimality. We expect these two parameters to be related, because larger noise strength requires stronger metabolic constant to prevent the activity of the network to be dominated by noise. We thus performed a 2-dimensional parameter search (Fig. 6I). As expected, the optima of the metabolic constant and the noise strength were positively correlated. A weaker noise required lower metabolic constant, and-vice-versa. While achieving maximal efficiency at non-zero levels of the metabolic cost and noise (see Fig. 6I) might seem counterintuitive, we speculate that such setting is optimal because some noise in the non-specific current prevents over-synchronization and over-regularity of firing that would harm efficiency, similarly to what was shown in previous works ^31,36,88^. In the presence of noise, a non-zero metabolic constant is needed to suppress inefficient spikes purely induced by noise that do not contribute to coding and increase the error. This gives rise to a form of stochastic resonance, where an optimal level of noise is helpful to detect the signal coming from the feedforward currents.

In summary, non-specific external currents derived in our optimal solution have a major effect on coding efficiency and on neural dynamics. In qualitative agreement with empirical measurements ^87,62^, our model predicts that more than half of the average distance between the resting potential and firing threshold is accounted for by non-specific synaptic currents. Similarly to previous theoretical work^31,36^, we find that some level of external noise, in the form of a random fluctuation of the non-specific synaptic current, is beneficial for network efficiency. This remains a prediction for experiments.

### Optimal ratio of E-I neuron numbers and of the mean I-I to E-I synaptic efficacy coincide with biophysical measurements

Next, we investigated how coding efficiency and neural dynamics depend on the ratio of the number of E and I neurons (*N* ^*E*^ : *N* ^*I*^ or E-I ratio) and on the relative synaptic strengths between E-I and I-I connections.

Efficiency objectives (Eq. 3) are based on population, rather than single-neuron activity. Our efficient E-I network thus realizes a computation of the target representation that is distributed across multiple neurons (Fig. 7A). Following previous reports ^37^, we predict that, if the number of neurons within the population decreases, neurons have to fire more spikes to achieve an optimal population readout because the task of tracking the target signal is distributed among fewer neurons. To test this prediction, we varied the number of I neurons while keeping the number of E neurons constant. As predicted, a decrease of the number of I neurons (and thus an increase in the ratio of the number of E to I neurons) caused a linear increase in the firing rate of I neurons, while the firing rate of E neurons stayed constant (Fig. 7B, top). However, the variability of spiking and the average synaptic inputs remained relatively constant in both cell types as we varied the E-I ratio (Fig. 7B, bottom, C), indicating a compensation for the change in the ratio of E-I neuron numbers through adjustment in the firing rates. These results are consistent with the observation in neuronal cultures of a linear change in the rate of postsynaptic events but unchanged postsynaptic current in either E and I neurons for variations in the E-I neuron number ratio^89^.

**Figure 7.**
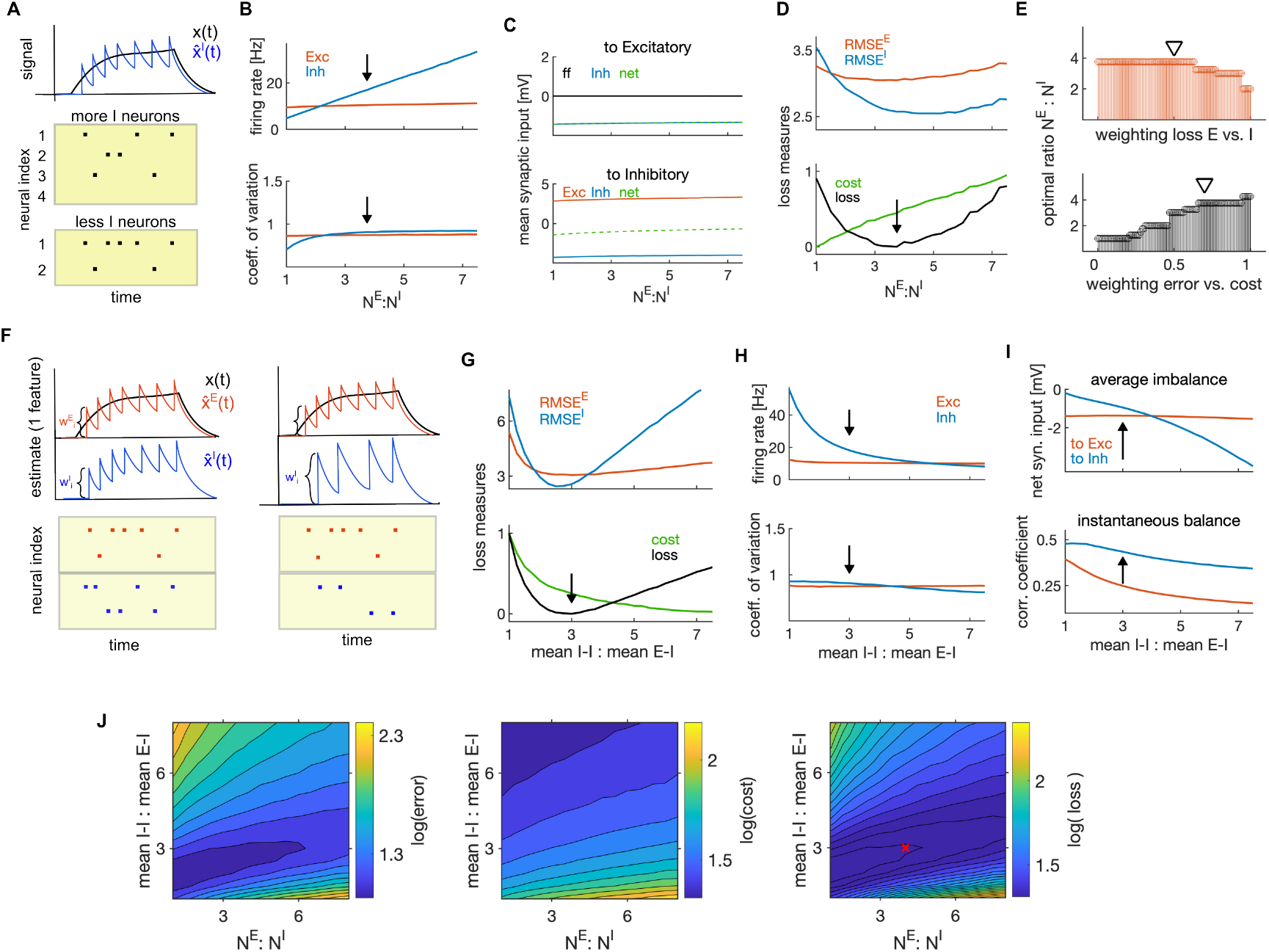
Optimal ratios of E-I neuron numbers and of mean I-I to E-I efficacy. **(A)** Schematic of the effect of changing the number of I neurons on firing rates of I neurons. As encoding of the stimulus is distributed among more I neurons, the number of spikes per I neuron decreases. **(B)** Average firing rate as a function of the ratio of the number of E to I neurons. Black arrow marks the optimal ratio. **(C)** Average net synaptic input in E neurons (top) and in I neurons (bottom). **(D)** Top: Encoding error (RMSE) of the E (red) and I (blue) estimates, as a function of the ratio of E-I neuron numbers. Bottom: Same as on top, showing the cost and the average loss. Black arrow shows the minimum of the loss, indicating the optimal parameter. **(E)** Top: Optimal ratio of the number of E to I neurons as a function of the weighting of the average loss of E and I cell type (using the weighting of the error and cost of 0.7 and 0.3, respectively). Bottom: Same as on top, measured as a function of the weighting of the error and the cost when computing the loss. (The weighting of the losses of E and I neurons is 0.5.) Black triangles mark weightings that we typically used. **(F)** Schematic of the readout of the spiking activity of E (red) and I population (blue) with equal amplitude of decoding weights (left) and with stronger decoding weight in I neuron (right). Stronger decoding weight in I neurons results in a stronger effect of spikes on the readout, leading to less spikes by the I population. **(G-H)** Same as in **D** and **B**, as a function of the ratio of mean I-I to E-I efficacy. **(I)** Average imbalance (top) and instantaneous balance (bottom) balance, as a function of the ratio of mean I-I to E-I efficacy. **(J)** The encoding error (RMSE; left) the metabolic cost (middle) and the average loss (right) as a function of the ratio of E-I neuron numbers and the ratio of mean I-I to E-I connectivity. The optimal ratios obtained with single parameter search (in **D** and **G**) are marked with a red cross. All statistical results were computed on 100 simulation trials of 1 second duration. For other parameters, see Table 1.

The ratio of the number of E to I neurons had a significant influence on coding efficiency. We found a unique minimum of the encoding error of each cell type, while the metabolic cost increased linearly with the ratio of the number of E and I neurons (Fig. 7D). Using the usual weighting *g*_*L*_ = 0.7, we found the optimal ratio of E to I neuron numbers to be in range observed experimentally in cortical circuits (Fig. 7D, bottom, black arrow, *N* ^*E*^ : *N* ^*I*^ = 3.75 : 1; ^90^). The optimal ratio depended on the weighting of the error with the cost, decreasing when increasing the cost of firing (Fig. 7E, bottom). Also the encoding error (RMSE) alone, without considering the metabolic cost, predicted optimal ratio of the number of E to I neurons within a plausible physiological range, *N* ^*E*^ : *N* ^*I*^ = [3.75 : 1, 5.25 : 1], with stronger weightings of the encoding error by I neurons predicting higher ratios (Fig. 7E, top).

Next, we investigated the impact of the strength of E and I synaptic efficacy (EPSPs and IPSPs). As evident from the expression for the population readouts (Eq. 2), the magnitude of tuning parameters (which are also decoding weights) determines the amplitude of jumps of the population readout caused by spikes (Fig. 7F). The larger these weights are, the larger is the impact of spikes on the population signals.

E and I synaptic efficacies depend on the tuning parameters. We parametrized the distribution of tuning parameters as uniform distributions centered at zero, but allowed the spread of distributions in E and I neurons (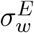 and 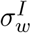) to vary across E and I cell type (Methods). In the optimally efficient network, as found analytically (Methods section “Dynamic equations for the membrane potentials”), the E-I connectivity is the transpose of the of the I-E connectivity, which implies that these connectivities are exactly balanced and have the same mean. We also showed analytically that by parametrizing tuning parameters with uniform distributions, the scaling of synaptic connectivity of E-I (equal to I-E) and I-I connectivity is controlled by the variance of tuning parameters of the pre and postsynaptic population as follows: 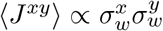. Using these insights, we were able to analytically evaluate the mean E-I and I-I synaptic efficacy (see Methods section “Parametrization of synaptic connectivity”).

We next searched for the optimal ratio of the mean I-I to E-I efficacy as the parameter that maximizes network efficiency. Network efficiency was maximized when such ratio was about 3 to 1 (Fig. 7G). Our results suggest the maximum E-I and I-E synaptic efficacy, averaged across neuronal pairs, of 0.75 mV, and the maximal I-I efficacy of 2.25 mV, values that are consistent with empirical measurements in the primary sensory cortex ^91,60,61^. The optimal ratio of mean I-I to E-I connectivity decreased when the error was weighted more with respect to the metabolic cost (Supplementary Fig. S4D).

Similarly to the ratio of E-I neuron numbers, a change in the ratio of mean E-I to I-E synaptic efficacy was compensated for by a change in firing rates, with stronger I-I synapses leading to a decrease in the firing rate of I neurons (Fig. 7H, top). Conversely, weakening the E-I (and I-E) synapses resulted in an increase in the firing rate in E neurons (Supplementary Fig. S4E-F). This is easily understood by considering that weakening the E-I and I-E synapses activates less strongly the lateral inhibition in E neurons (Fig. 3) and thus leads to an increase in the firing rate of E neurons. We also found that single neuron variability remained almost unchanged when varying the ratio of mean I-I to E-I efficacy (Fig. 7H, bottom) and the optimal ratio yielded optimal levels of average and instantaneous balance of synaptic inputs, as found previously (Fig. 7I). The instantaneous balance monotonically decreased with increasing ratio of I-I to E-I efficacy (Fig. 7I, bottom, Supplementary Fig. S4G).

Further, we tested the co-dependency of network optimality on the above two ratios with a 2-dimensional parameter search. We expected a positive correlation of network performance as a function of these two parameters, because both of them regulate the level of instantaneous E-I balance in the network. We found that the lower ratio of E-I neuron numbers indeed predicts a lower ratio of the mean I-I to E-I connectivity (Fig. 7J). This is because fewer E neurons bring less excitation in the network, thus requiring less inhibition to achieve optimal levels of instantaneous balance. The co-dependency of the two parameters in affecting network optimality might be informative as to why E-I neuron number ratios may vary across species (for example, it is reported to be 2:1 in human cortex ^92^ and 4:1 in mouse cortex). Our model predicts that lower E-I neuron number ratios require weaker mean I-I to E-I connectivity.

In summary, our analysis suggests that optimal coding efficiency is achieved with more E neurons than I neurons and with mean I-I synaptic efficacy stronger than the E-I and I-E efficacy, and that these two parameters are positively correlated. Optimal ratios of E to I neurons and of connection strengths are broadly consistent with empirical measurements of these parameters in biological networks. The optimal network has less I than E neurons, but the impact of spikes of I neurons on the population readout is stronger, also suggesting that spikes of I neurons convey more information.

### Dependence of efficient coding and neural dynamics on the stimulus statistics

We further investigated how the network’s behavior depends on the timescales of the input stimulus features. We manipulated the stimulus timescales by changing the time constants of *M* = 3 OU processes. The network efficiently encoded stimulus features when their time constants varied between 1 and 200 ms, with stable encoding error, metabolic cost (Fig. 8A) and neural dynamics (Supplementary Fig. S5 A-B). To examine if the network can efficiently encode also stimuli that evolve on different timescales, we tested its performance in response to *M* = 3 input variables, each with a different timescale. We kept the timescale of the first variable constant at 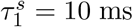, while we varied the time constants of the other two keeping the time constant of the third twice as long as that of the second. We found excellent performance of the network in response to such stimuli that was stable across timescales (Fig. 8B). The prediction that the network can encode information effectively over a wide range of time scales can be tested experimentally, by measuring the sensory information encoded by the activity of a set of neurons while varying the sensory stimulus timescales over a wide range.

**Figure 8.**
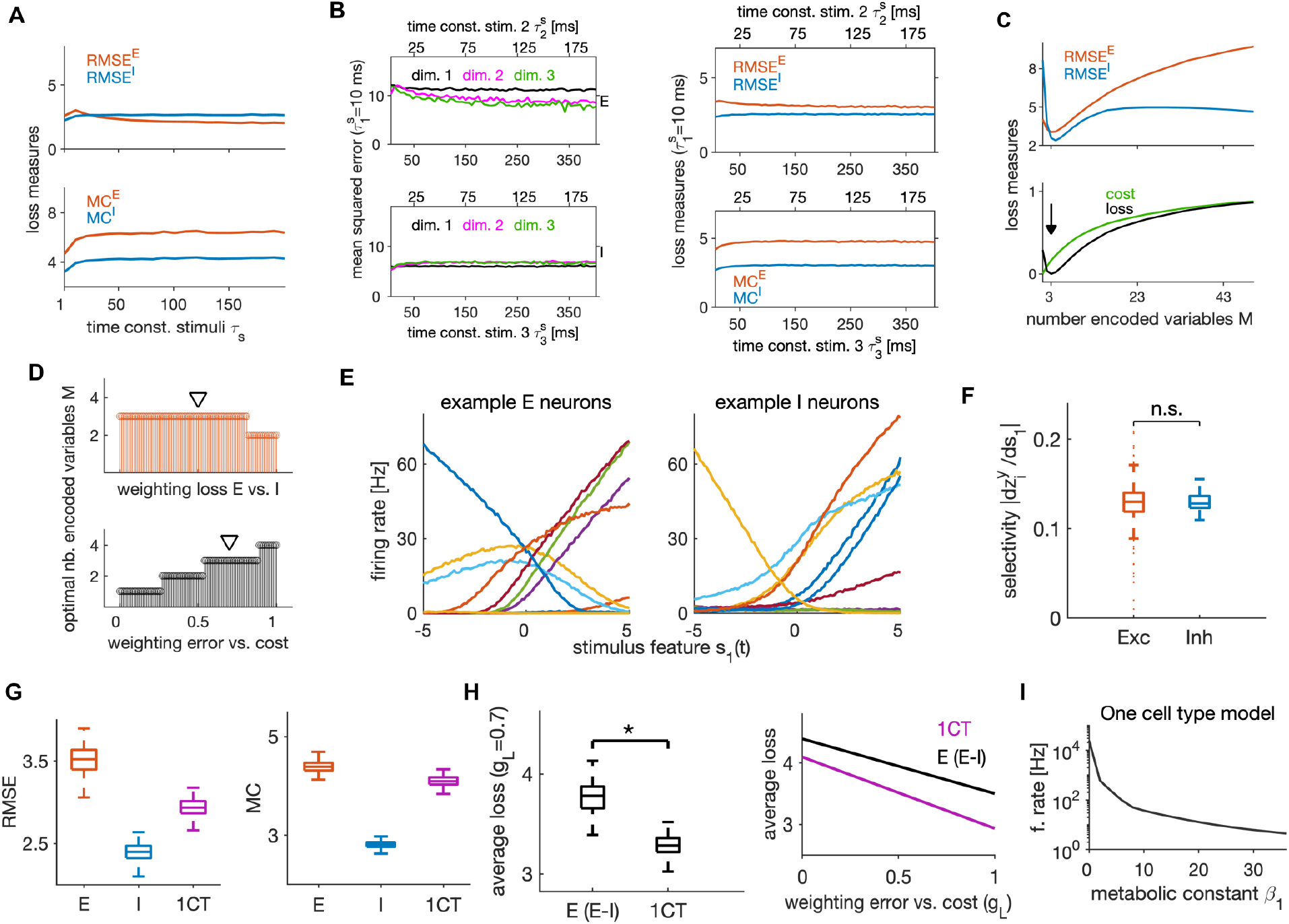
Dependence of efficient coding and neural dynamics on stimulus parameters and comparison of E-I versus one cell type model architecture. **(A)** Top: Root mean squared error (RMSE) of E estimates (red) and I estimates (blue), as a function of the time constant (in ms) of stimulus features. The time constant *τ*_*s*_ is the same for all stimulus features. Bottom: Same as on top, showing the metabolic cost (MC) of E and I cell type. **(B)** Left: Mean squared error between the targets and their estimates for every stimulus feature (marked as dimensions), as a function of time constants of OU stimuli in E population (top) and in I population (bottom). In the first dimension, the stimulus feature has a time constant fixed at 10 ms, while the second and third feature increase their time constants from left to right. The time constant of the third stimulus feature (x-axis on the bottom) is the double of the time constant of the second stimulus feature (x-axis on top). Right: Same as on the left, showing the RMSE that was averaged across stimulus features (top), and the metabolic cost (bottom) in E (red) and I (blue) populations. **(C)** Top: Same as in **A** top, measured as a function of the number of stimulus features *M*. Bottom: Normalized cost and the average loss as a function of the number of stimulus features. Black arrow marks the minimum loss and the optimal parameter *M*. **(D)** Top: Optimal number of encoded variables (stimulus features) as a function of weighting of the losses of E and I population. The weighting of the error with the cost is 0.7. Bottom: Same as on top, as a function of the weighting of the error with the cost and with equal weighting of losses of E and I populations. **(E)** Tuning curves of 10 example E (left) and I neurons (right). We computed tuning curves using *M* =3 stimulus features that were constant over time. We varied the amplitude of the first stimulus feature *s*_1_, while two other stimulus features were kept fixed. **(F)** Distribution of the selectivity index across E (red) and I neurons (blue). **(G)** Root mean squared error (left) and metabolic cost (right) in E and I populations in the E-I model and in the 1CT model. The distribution is across 100 simulation trials. **(H)** Left: Average loss in the E population of the E-I model and of the 1CT model. The distribution is across 100 simulation trials. Right: Average loss in the E population of the E-I models and in the 1CT model as a function of the weighting *g*_*L*_, averaged across trials. **(I)** Firing rate in the 1CT model as a function of the metabolic constant. All statistical results were computed on 100 simulation trials of 1 second duration. For other parameters of the E-I model see Table 1, and for the 1CT model see Supplementary Table S2

We next examined network performance while varying the timescale of targets *τ*_*x*_ (see Eq. 1). Because we assumed that the target time constants equal the membrane time constant of E and I neurons (*τ*_*x*_ = *τ* ^*E*^ = *τ* ^*I*^ = *τ*), it is not surprising that the best performance was achieved when these time constants were similar (Supplementary Fig. S5C). Firing rates, firing variability and the average and instantaneous balance did not change appreciably with this time constant (Supplementary Fig. S5D-E).

Next, we tested how the network’s behavior changed when we varied the number of stimulus features *M*. Because all other parameters were optimized using *M* = 3, the encoding error of E (RMSE^*E*^) and I neurons (RMSE^*I*^) achieved a minimum around this value (Fig. 8C, top). The metabolic cost increased monotonically with *M* (Fig. 8C, bottom). The number of features that optimized network efficiency (and minimized the average loss) depended on *g*_*L*_, with stronger penalty of firing yielding a smaller optimal number of features. Increasing *M* beyond the optimal number resulted in a gentle monotonic increase in firing rates for both E and I neurons, and it increased the average E-I balance and weakened the instantaneous balance (Supplementary Fig. S5F-G).

We next characterized the tuning and the stimulus selectivity of E and I neurons. E neurons receive a feedforward current, which is expected to make them stimulus-selective, while I neurons receive synaptic inputs from E neurons through dense E-I connectivity. We measured stimulus tuning by computing tuning curves for each neuron in response to *M* =3 constant stimulus features (see Methods). Similarly to previous work^37^, tuning curves of both E and I neurons were strongly heterogeneous (Fig. 8E). We tested if the selectivity differs across E and I cell types. We computed a selectivity index for each neuron as the stimulus-response gain (average change in the firing rate in response to a small change in the stimulus divided by the stimulus change size, see Methods), and found that E and I neurons had similar mean stimulus selectivity (*p* = 0.418, two-tailed t-test; Fig. 8F). Thus, I neurons, despite not receiving direct feedforward inputs and acquiring stimulus selectivity only through structured E-I connections, are tuned to the input stimuli as strongly as the E neurons.

### Comparison of E-I and one cell type model architecture for coding efficiency and robustness

Neurons in the brain are either excitatory or inhibitory. To understand how differentiating E and I neurons benefits efficient coding, we compared the properties of our efficient E-I network with an efficient network with a single cell type (1CT). The 1CT model can be seen as a simplification of the E-I model (see Supplementary Text 1) and has been derived and analyzed in previous studies ^29,28,36,33,93,43^. We compared the average encoding error (RMSE), the average metabolic cost (MC), and the average loss (see Supplementary Text 3) of the E-I model against the one cell type (1CT) model. Compared to the 1CT model, the E-I model exhibited a higher encoding error and metabolic cost in the E population, but a lower encoding error and metabolic cost in the I population (Fig. 8G). The 1CT model can perform similar computations as the E-I network. Instead of an E neuron directly providing lateral inhibition to its neighbor (Supplementary Fig. S1A-C), it goes through an interneuron in the E-I model (Fig. 1A(i) and B). Because the E population of the E-I model and the 1CT model perform a similar computation, we compared the efficiency of the E population of the E-I model with the 1CT model. We found that the 1CT model is slightly more efficient than the E population of the E-I model, consistently for different weightings of the error with the cost (Fig. 8H).

We further compared the robustness of firing rates to changes in the metabolic constant of the two models. Consistently with previous studies ^36,35^, firing rates in the 1CT model were highly sensitive to variations in the metabolic constant (Fig. 8I, note the logarithmic scale on the y-axis), with a superexponential growth of the firing rate with the inverse of the metabolic constant in regimes with metabolic cost lower than optimal. This is in contrast to the E-I model, whose firing rates exhibited lower sensitivity to the metabolic constant, and never exceeded physiological limits (Fig. 6C, top). Because our E-I model does not incorporate a saturating input-output function that constrains the range of firing as in^34^, the ability of the E-I model to maintain firing rates within biologically plausible limits emerges as a highly desirable dynamic property. One reason for higher stability of our E-I model compared to the 1CT model is that the delay of the lateral inhibition in the E-I model is twice that of the 1CT model (because in the E-I model, the lateral inhibition travels through an additional synapse). A second reason is that the recurrent connectivity of the 1CT model has exactly the same amount of average excitation and inhibition, while the E-I model is inhibition-dominated, which makes the E-I model more stable.

In summary, although the optimal E-I model is slightly less efficient than the optimal 1CT model, it does not enter into states of physiologically unrealistic firing rates when the metabolic constant is lower than the optimal one.

## Discussion

We analyzed the structural, dynamical and coding properties that emerge in networks of spiking neurons that implement efficient coding. We demonstrated that efficient recurrent E-I networks form highly accurate representations of stimulus features with biologically plausible parameters, biologically plausible neural dynamics, instantaneous E-I balance and like-to-like connectivity structure leading to lateral inhibition. The network can implement efficient coding with stimulus features varying over a wide range of timescales and when encoding even multiple such features. Here we discuss the implications of these findings.

By a systematic study of the model, we determined the model parameters that optimize network efficiency. The optimal parameters (including the ratio between the number of E and I neurons, the ratio of I-I to E-I synaptic efficacy and parameters of non-specific currents) were consistent with parameters measured empirically in cortical circuits, and generated plausible spiking dynamics. This result lends credibility to the hypothesis that cortical networks might be designed for efficient coding and may operate close to optimal efficiency, as well as provides a solid intuition about what specific parameter ranges (e.g. higher numbers of E than I neurons) may be good for. With moderate deviations from the optimal parameters, efficient networks still exhibited realistic dynamics and close-to-efficient coding, suggesting that the optimal operational point of such networks is relatively robust. We also found that optimally efficient analytical solution derives generalized LIF (gLIF) equations for neuron models ^38^. While gLIF ^83,41^ and LIF ^77,78^ models are reasonably biologically plausible and are widely used to model and study spiking neural network dynamics, it was unclear how their parameters affect network-level information coding. Our study provides a principled way to determine uniquely the parameter values of gLIF networks that are optimal for efficient information encoding. Studying the dynamics of gLIF networks with such optimal parameters thus provides a direct link between optimal coding and neural dynamics. Moreover, our formalism provides a framework for the optimization of neural parameters that can in principles be used not only for neural network models that study brain function but also for the design of artificial neuromorphic circuits that perform information coding computations ^94,95^.

Our model generates a number of insights about the role of structured connectivity in efficient information processing. A first insight is that I neurons develop stimulus feature selectivity because of the structured recurrent connectivity. While in visual cortex of naive animals, I neurons are typically reported to be less tuned than E neurons^96,97^ (but see ^98^), recent studies in association areas of well task trained animals consistently find I neurons as strongly tuned as E neurons ^99,100^. A second insight is that a network with structured connectivity shows stronger average and instantaneous E-I balance, as well as significantly lower variance of membrane potentials compared to an equivalent network with randomly organized connections. This implies that the connectivity structure is not only crucial for coding efficiency, but also influences the dynamical regime of the network. A third insight is that the structured network exhibits both lower encoding error and lower firing rates compared to unstructured networks, thus achieving higher efficiency. Our analysis of the effective connectivity created by the efficient connectivity structure shows that this structure sharpens stimulus representations, reduces redundancy and increases metabolic efficiency by implementing feature-specific competition, that is a negative effective connectivity between E neurons with similar stimulus tuning, as proposed by recent theories^30^ and experiments ^58,59^ of computations in visual cortex.

Our model gives insights on what would be minimal requirements for a biological network to implement efficient coding. The network has to have structured E-I and I-E connectivity and weak and short-lasting or no spike-triggered adaptation. Further, at least half of the distance between the resting potential and the threshold should be provided by a stochastic external current that is unrelated to the feedforward stimuli. Finally, the network should have a ratio of E to I neuron numbers in the range of about 2:1 to 4:1 and the ratio of average I-I to E-I connectivity in the range of about 2:1 to 3:1, with smaller E-I neuron number ratios implying smaller average I-I to E-I connectivity ratios.

Our study gives insights into how structured connectivity between E and I neurons affects the dynamics of E-I balancing and how this relates to information coding. Previous work^32^ proposed that the E-I balance in efficient spiking networks operates on a finer time scale than in classical balanced E-I networks with random connectivity^78^. However, theoretical attempts to determine the levels of instantaneous E-I balance that are optimal for coding are rare^101^. Consistent with the general idea put forth in ^32,31,53^, we here showed that moderate levels of E-I balance are optimal for coding, and that too strong levels of instantaneous E-I balance are detrimental to coding efficiency. Our results predict that structured E-I-E connectivity is necessary for optimal levels of instantaneous E-I balance. Finally, the E-I-E structured connectivity that we derived supports optimal levels of instantaneous E-I balance and causes desynchronization of the spiking output. Such intrinsically generated desynchronization is a desirable network property that in previously proposed models could only be achieved by the less plausible addition of strong noise to each neuron^31,35^.

Our result that network efficiency depends gently on the number of neurons is consistent with previous findings that demonstrated robustness of efficient networks to neuronal loss^37^ and robustness of efficient spiking to the number of neurons^80^. Building on these studies, we additionally documented how the optimal ratio of the number of E to I neurons relates to the optimal ratio of average I-I to E-I connectivity. In particular, our analysis predicts that the optima of these two ratios are positively correlated. This might give insights into the diversity of ratios of E-I neuron number ratios observed across species ^92^.

We found that our efficient network, optimizing the representation of a leaky integration of stimulus features, does not require recurrent E-E connections. This is compatible with the relatively sparse levels of recurrent E-E connections in primary visual cortex ^102^, with the majority of E-E synapses suggested to be long-range^103^. Nevertheless, a limitation of our study is that it did not investigate the computations that could be made by E-E connections. Future studies could address the role of recurrent excitatory synapses that implement efficient coding computations beyond leaky integration, such as linear^38^ or non-linear mixing of stimulus features ^93^. Investigating such networks would also allow addressing whether biologically plausible efficient networks exhibit criticality, as suggested by ^104^.

A more realistic mapping of efficient coding onto biological networks would also entail including multiple types of inhibitory neurons ^105^, which could provide additional insights into how interneuron diversity serves information coding. Further limitations of our study to be addressed in future work include a more realistic implementation of the feedforward current. In our implementation, the feedforward current is simply a sum of uncorrelated stimulus features. However, in biological circuits, the feedforward input is a series of complex synaptic inputs from upstream circuits. A more detailed implementation of feedforward inputs, coupled with recurrent E-E synapses, might influence the levels of the instantaneous balance, in particular in E neurons, and have an impact on network efficiency. Moreover, we here did not explore cases where the same stimulus feature has multiple time scales. Finally, we note that efficient encoding might be the primary normative objective in sensory areas, while areas supporting high-level cognitive tasks might include other computational objectives, such as efficient transmission of information downstream to generate reliable behavioral outputs ^106,107,108,25,109^. It would thus be important to understand how networks could simultaneously optimize or trade off different objectives.

## Acknowledgements

V.K. and T.S. thank Tatiana Engel for her contribution to the discussion of results and for her comments on an earlier version of the manuscript. This project was supported by funding from Technische Universität Berlin (“Equal Opportunity Program” to VK), by Internal Research Funding of Technische Universität Berlin (to TS), by NIH Brain Initiative (grants U19 NS107464, R01 NS109961, R01 NS108410 to SP), and the Simons Foundation for Autism Research Initiative (SFARI; grant 982347 to SP), and by the European Union’s Horizon 2023 research and innovation program under the Marie Sklodowska-Curie Actions (grant 101152984 ASTRONET to SBM).

## Code availability

The complete computer code for reproducing the results can be downloaded anonymously from a public GitHub repository https://github.com/VeronikaKoren/efficient_EI and has the associated DOI: https://doi.org/10.5281/zenodo.14628524.

## Methods

### Overview of the current approach and of differences with previous approaches

In the following, we present a detailed derivation of the E-I spiking network implementing the efficient coding principle. The analytical derivation is based on previous works on efficient coding with spikes ^28,36^, and in particular on our recent work ^38^. While these previous works analytically derived feedforward and recurrent transmembrane currents in leaky integrate-and fire neuron models, they did not contain any synaptic currents unrelated to feedforward and recurrent processing. Non-specific synaptic currents were suggested to be important for an accurate description of coding and dynamics in cortical networks ^62^. In the model derivation that follows, we also derived non-specific external current from efficiency objectives.

Moreover, we here revisited the derivation of physical units in efficient spiking networks. We built on a previous work ^28^ that assigned physical units to mathematical expressions that correspond to membrane potentials, firing thresholds, etc. Here, we instead assigned physical units to the computational variables such as the target signals and the population readouts, and then derived units of the membrane potentials and firing thresholds.

With this model, we aim to describe neural dynamics and computation in early sensory cortices such as the primary visual cortex in rodents, even though many principles of the model developed here could be relevant throughout the brain.

### Introducing variables of the model

We consider two types of neurons, excitatory neurons *E* and inhibitory neurons *I*. We denote as *N* ^*E*^ and *N* ^*I*^ the number of *E*-cells and *I*-cells, respectively. The spike train of neuron *i* of type *y* ∈ {*E, I*}, *i* = 1, 2, …, *N* ^*y*^, is defined as a sum of Dirac delta functions,

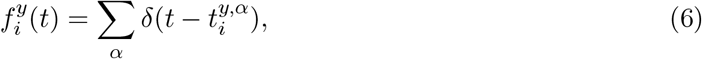

where 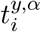 is the time of the *α*-th spike of that neuron, defined as a time point at which the membrane potential of neuron *i* crosses the firing threshold.

We define the readout of the spiking activity of neuron *i* of type *y* (in the following, “single neuron readout”) as a leaky integration of its spike train,

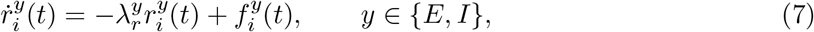

with λ_*r*_ denoting the inverse time constant. This way, the quantity 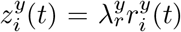 represents an estimate of the instantaneous firing rate of neuron *i*.

We denote as ***s***(*t*) := [*s*_1_(*t*), …, *s*_*M*_ (*t*)]^⊤^ the set of *M* dynamical features of the external stimulus (in the following, stimulus features) which are transmitted to the network through a feedforward sensory pathway. The stimulus features have the unit of the square root of millivolt, 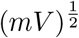. The target signal is then obtained through a leaky integration of the feedforward variable, ***s***(*t*) ^29^, with inverse time constant λ, as

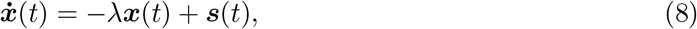

with ***x***(*t*) := [*x*_1_(*t*), …, *x*_*M*_ (*t*)]^⊤^ the vector of *M* target signals. Furthermore, we define a linear population readout of the spiking activity of E and I neurons

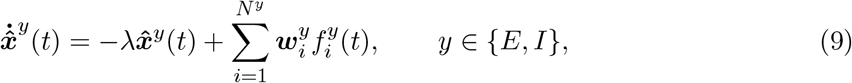

with 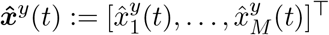 the vector of estimates of cell type *y* and 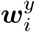 in units of 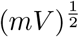. Here, each neuron *i* of type *y* is associated with a vector 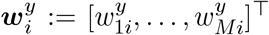 of *M* tuning parameters representing the decoding weight of neuron *i* with respect to the *M* population readouts in Eq. 9. These decoding vectors can be combined in the *M* × *N* ^*y*^ matrix 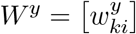. The rows of this matrix define the patterns of decoding weights 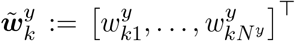 for each signal dimension *k* = 1, …, *M*.

### Loss functions

We assume that the activity of a population *y* ∈ {*E, I*} is set so as to minimize a time-dependent encoding error and a time-dependent metabolic cost:

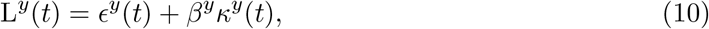

with *β*^*y*^ *>* 0 in units of mV the Lagrange multiplier which controls the weight of the metabolic cost relative to the encoding error. The time-dependent encoding error is defined as the squared distance between the targets and their estimates, and the role of estimates is assigned to the population readouts 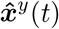. In E neurons, the targets are defined as the target signals ***x***(*t*), and their estimators are the population readouts of the spiking activity of E neurons, 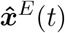. In I neurons, the targets are defined as the population readouts of E neurons 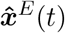 and their estimators are the population readouts of I neurons 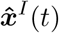. Furthermore, the time-dependent metabolic cost is proportional to the squared estimate of the instantaneous firing rate, summed across neurons from the same population. Following these assumptions, we define the variables of loss functions in Eq. 10 as

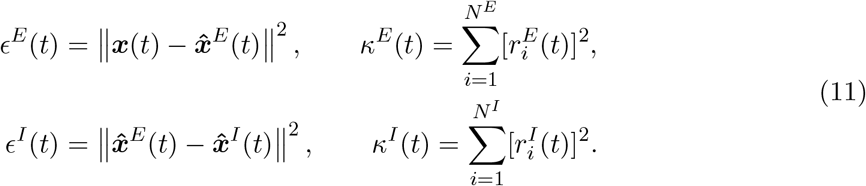

We use a quadratic metabolic cost because it promotes the distribution of spiking across neurons ^28^. In particular, the loss function of I neurons, L^*I*^(*t*) implies the relevance of the approximation: 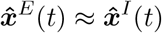 (see *ϵ*^*I*^ in the Eq. 11), which will be used in what follows.

### When shall a neuron spike?

We minimize the loss function by positing that neuron *i* of type *y* ∈ {*E, I*} emits a spike as soon as its spike decreases the loss function of its population *y* in the immediate future^38^. We also define *t*^−^ and *t*^+^ as the left- and right-sided limits of a spike time 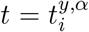, respectively. Thus, at the spike time, the following jump condition must hold:

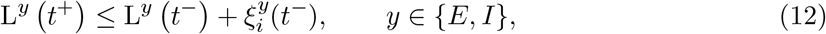

with 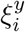 in units of mV. Here, the arguments *t*^−^ and *t*^+^ denote the left- and right-sided limits of the respected functions at time *t*. Furthermore, we added a noise term on the right-hand side of the Eq. (12) in order to consider the stochastic nature of spike generation in biological networks ^54^. A convenient choice for the noise 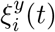 is the Ornstein-Uhlenbeck process obeying

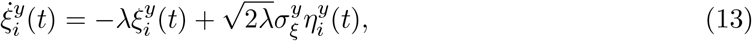

where 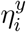 is a Gaussian white noise with auto-covariance function ⟨*η*_*i*_(*t*)*η*_*j*_(*t*^′^)⟩ = *δ*_*ij*_*δ*(*t* − *t*^′^). The process 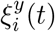 has zero mean and auto-covariance function 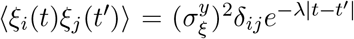, with 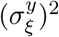 the variance of the noise.

By applying the condition for spiking in Eq. (12) using *y* = *E* and *y* = *I*, respectively, we get

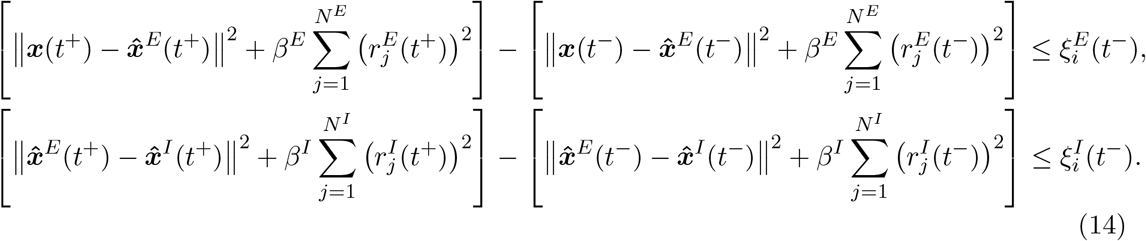

According to the definitions in Eqs. (7) and (9), if neuron *i* fires a spike at time 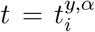, it causes a jump of its own filtered spike train (but not of other neurons *j* ≠ *i*), as well as of the population readout of the population it belongs to. Therefore, when neuron *i* fires a spike, we have for a given neuron *j* and a given population readout *k*:

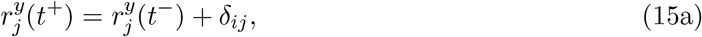

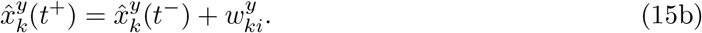

By inserting Eq. (15a)-(15b) in Eq. (12), we find that neuron *i* of type *y* should fire a spike if the following condition holds:

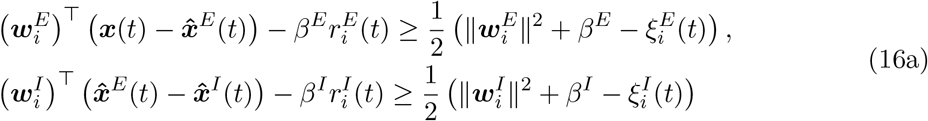

with 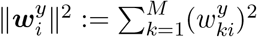 the squared length of the tuning vector of neuron *i* of type *y*. These equations tell us when the neuron *i* of type *E* and *I*, respectively, emits a spike, and are similar to the ones derived in previous works ^38,28^. In addition to what has been found in these previous works, we here also find that each term on the left- and right-hand side in the Eq 16a has the physical units of millivolts.

We note that the expression derived from the minimization of the loss function of E neurons in the top row of Eq. (16a) is independent of the activity of I neurons, and would thus lead to the E population being unconnected with the I population. In order to derive a recurrently connected E-I network, the activity of E neurons must depend on the activity of I neurons. We impose this property by using the approximation of estimates that holds under the assumption of efficient coding in I neurons (see *ϵ*^*I*^ in the Eq. 11), 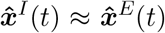. This yields the following conditions:

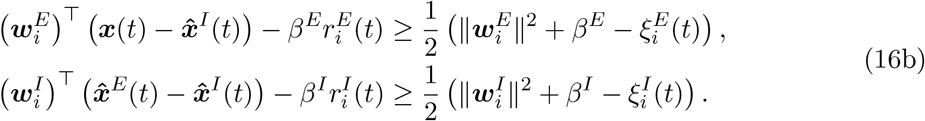

We now define new variables 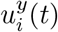 and 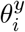 as proportional to the left- and the right-hand side of these expressions,

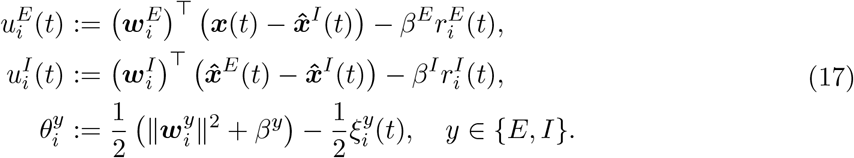

The variables 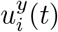 and 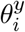 are interpreted as the membrane potential and the firing threshold of neuron *i* of cell type *y* ∈ {*E, I*}.

### Dynamic equations for the membrane potentials

In this section we develop the exact dynamic equations of the membrane potentials 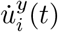 for *y* ∈ {*E, I*} according to the efficient coding assumption. We rewrite Eq. (9) in vector notation as

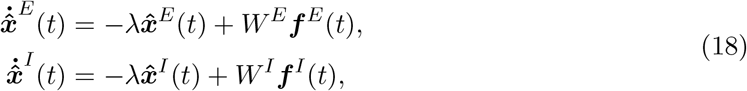

with 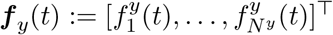 the vector of spike trains for *N* ^*y*^ neurons of cell type *y* ∈ {*E, I*}. In the case of E neurons, the time-derivative of the membrane potential 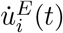 in Eq. (17), is obtained as

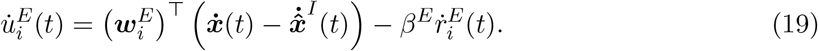

By inserting the dynamic equations of the target signal 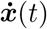, its estimate 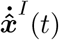 (Eq. 18) and of the single neuron readout 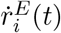 (Eq. 7 in the case *y* = *E*), we get

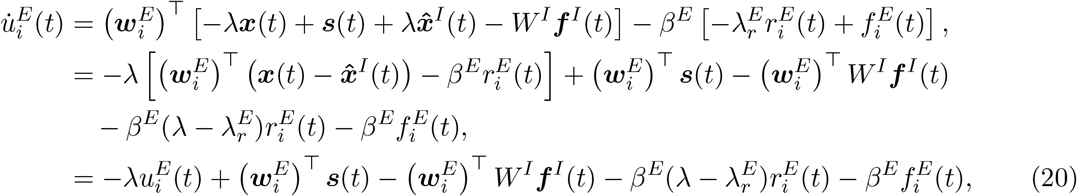

where in the last line we used the definition of 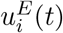 from the Eq. (17).

In the case of I neurons, the time derivative of the membrane potential 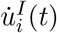 in Eq. (17) is

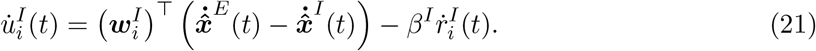

By inserting the dynamic equations of the population readouts of E neurons 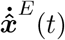 and of the I neurons 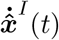 (Eq. 18) and of the single neuron readout 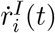 (Eq. 7 in the case *y* = *I*), we get

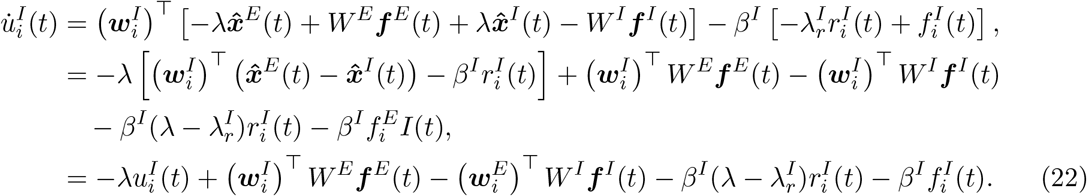

where in the last line we used the definition of 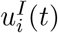 from Eq. (17).

### Leaky integrate-and-fire neurons

The terms on the right-hand-side in Eqs. (20) and (22) can be interpreted as transmembrane currents. The last term in these equations, 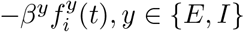, can be interpreted as a current instantaneously resetting the membrane potential upon reaching the firing threshold^28^. Indeed, when the membrane potential reaches the threshold, it triggers a spike and causes a jump of the membrane potential by an amount −*β*^*y*^; this realizes resetting of the membrane potential which is equivalent to the resetting rule of integrate-and-fire neurons^64,66^. Thus, by taking into account the resetting mechanism and defining the time constants of population and single neuron readout *τ* := λ^−1^ and 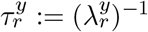, we can rewrite Eqs. (20) and (22) as a leaky integrate-and-fire neuron model,

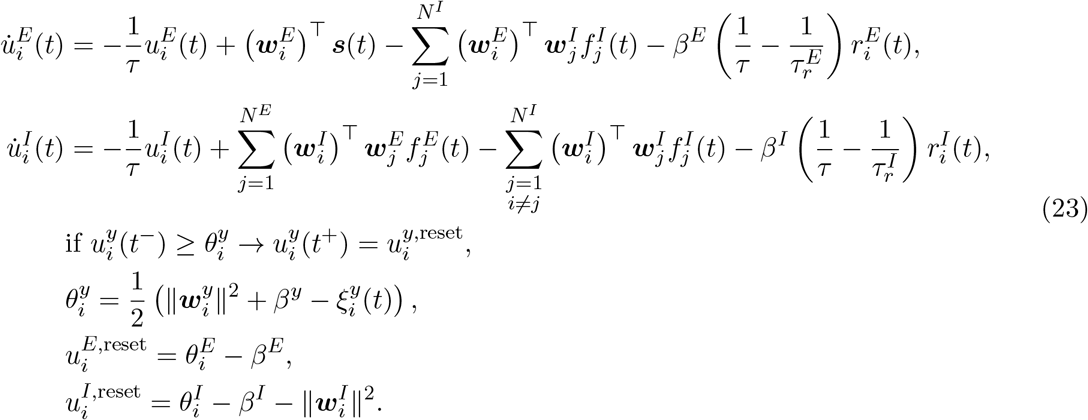

In the Eq. 23 we wrote explicitly the terms 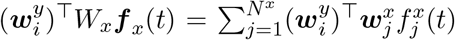, which correspond to the synaptic projections of *N* ^*x*^ presynaptic neurons of type *x* to the postsynaptic neuron *i* of type *y*, with the quantity 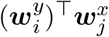 denoting the synaptic weight. We note that, in the case of I neurons, the element with *j* = *i* describes an autapse, i.e., a projection of a neuron with itself; this term is equal to 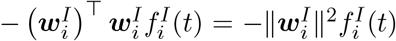, and thus contributes to the resetting of the neuron *i*.

### Imposing Dale’s principle on synaptic connectivity

We now examine the synaptic terms in Eq. (23). As a first remark, we see that synaptic weights depend on tuning parameters 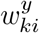. For the sake of generality we drew tuning parameters 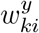 from a normal distribution with vanishing mean, which yielded both positive and negative values of 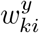. This has the desirable consequence that a spike of a neuron with a positive tuning parameter in signal dimension *k*, 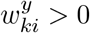 pulls the estimate, 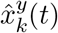, up, while a spike of a neuron with 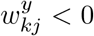 pulls the estimate down, allowing population readouts to track both positive and negative fluctuations of the target signal on a fast time scale.

Another consequence of synaptic connectivity in the Eq. (23) is that the synaptic weight between a presynaptic neuron *j* of type *x* and a postsynaptic neuron *i* of type *y* is symmetric and depends on the similarity of tuning vectors of the presynaptic and the postsynaptic neuron: 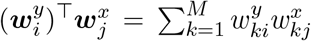. The sign of this scalar product is positive between neurons with similar tuning and negative between neurons with different tuning (and zero when the two tuning vectors are orthogonal). Thus, for a presynaptic neuron *j* of type *x*, the synaptic weights of its outgoing connections can be both positive and negative, because some of its postsynaptic neurons have similar tuning to the neuron *j* while others have different tuning. This is inconsistent with Dale’s principle^110^, which postulates that a particular neuron can only have one type of effect on postsynaptic neurons (excitatory or inhibitory), but never both. To impose this constraint in our model, we set synaptic weights between neurons with different tuning 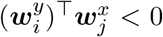 to zero. To this end, we define the rectified connectivity matrices,

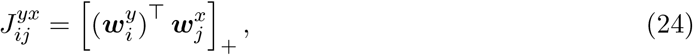

with (*x, y*) ∈ {(*E, I*), (*I, I*), (*I, E*)} and [*a*]_+_ ≡ max(0, *a*) a rectified linear function. Note that there are no direct synaptic connections between E neurons. Since the elements of the matrix *J* ^*yx*^ are all non-negative, it is the sign in front of the synaptic term in the Eq. (23) that determines the sign of the synaptic current between neurons *i* and *j*. The synaptic current is excitatory if the sign is positive, and inhibitory if the sign is negative.

It is also interesting to note that rectification affects the rank of connectivity matrices. Without rectification, the product in Eq. (24) yields a connectivity matrix with rank smaller or equal to the number of input features to the network, *M*, similarly as in previous works ^29,37,93^. Since typically the number of input features is much smaller than the number of neurons, i.e., *M <*< *N* ^*y*^, this would give a low-rank connectivity matrix. However, rectification in Eq. (24), necessary to ensure Dale’s principle in presence of positive and negative tuning parameters, typically results in a substantial increase of the rank of the connectivity matrix.

Using the synaptic connectivity defined in Eq. (24), we rewrite the network dynamics from Eq. (23) as:

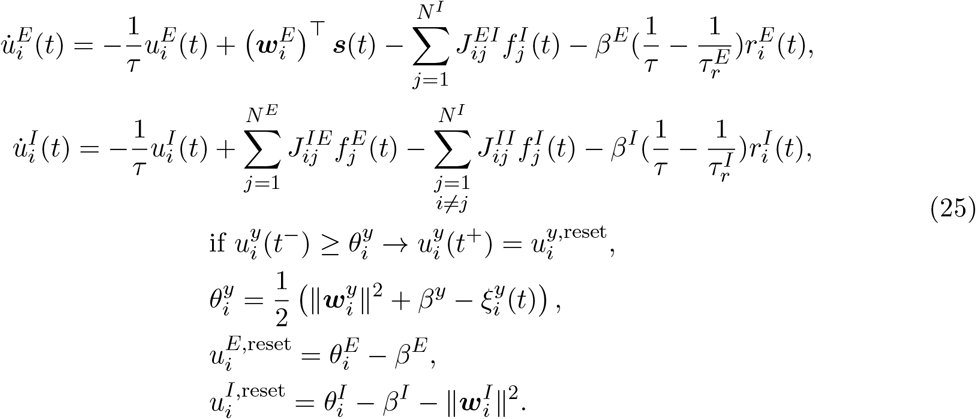

These equations express the neural dynamics which minimizes the loss functions (Eq. (10)) in terms of a generalized leaky integrate-and-fire model with E and I cell types, and are consistent with Dale’s principle.

In principle, it is possible to use the same strategy as for the E-I network to enforce Dale’s principle in model with one cell type (introduced by^28^). To do so, we constrained the recurrent connectivity of the model with a single cell type from ^36^ by keeping only connections between neurons with similar tuning vectors and setting other connections to 0 (see Supplementary Text 1). This led to a network of only inhibitory neurons, a type of network model which is less relevant for the description of biological networks.

### Model with resting potential and an external current

In the model given by the Eq. (25) the resting potential is equal to zero. In order to account for biophysical values of the resting potential and to introduce an implementation of the metabolic constant that is consistent with neurobiology, we add a constant value to the dynamical equations of the membrane potentials 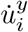, the firing thresholds 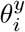 and the reset potentials 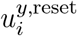. This does not change the spiking dynamics of the model, as what matters to correctly infer the efficient spiking times of neurons is the distance between the membrane potential and the threshold. Furthermore, in the same equations, the role of the metabolic constant *β*^*y*^ as a biophysical quantity is questionable. The metabolic constant *β*^*y*^ is an important parameter that weights the metabolic cost over the encoding error in the objective functions (Eq. 10). On the level of computational objectives, the metabolic constant naturally controls firing rates, as it allows the network to fire more or less spikes to correct for a certain encoding error. A flexible control of the firing rates is a desirable property, as gives the possibility to potentially capture different dynamical regimes of efficient spiking networks ^36^. In the spiking model we developed thus far (Eq. 25), similarly to previous efficient spiking models ^36,33^, the metabolic constant *β*^*y*^ controls the firing threshold. In neurobiology, however, strong changes to the firing threshold that would reflect metabolic constraints of the network are not plausible. We thus searched for an implementation of the metabolic constant *β*^*y*^ that is consistent with neurobiology.

The condition for threshold crossing of the neuron *i* can be written by Eq. (25) as

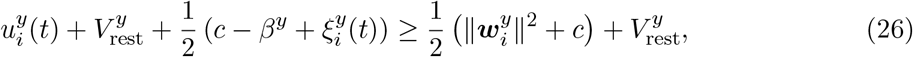

with *c* an arbitrary constant in units of millivolts. In Eq. (26) we added a constant *c*/2 and a resting potential 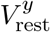 on the left- and right-hand side of the firing rule. Moreover, we shifted the noise and the dependency on the parameter *β* from the firing threshold to the membrane potential. Thus, we assumed that the firing threshold is independent of the metabolic constant and the noise, and we instead assumed the dependence on the metabolic constant and noise in the membrane potentials.

We now define new variables for *y* ∈ {*E, I*}:

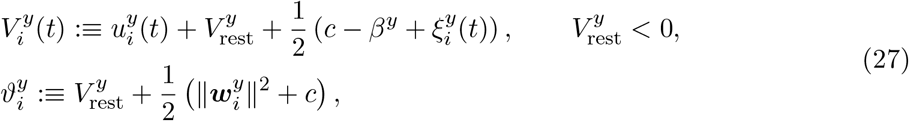

and rewrite the model in Eq. 25 in these new variables

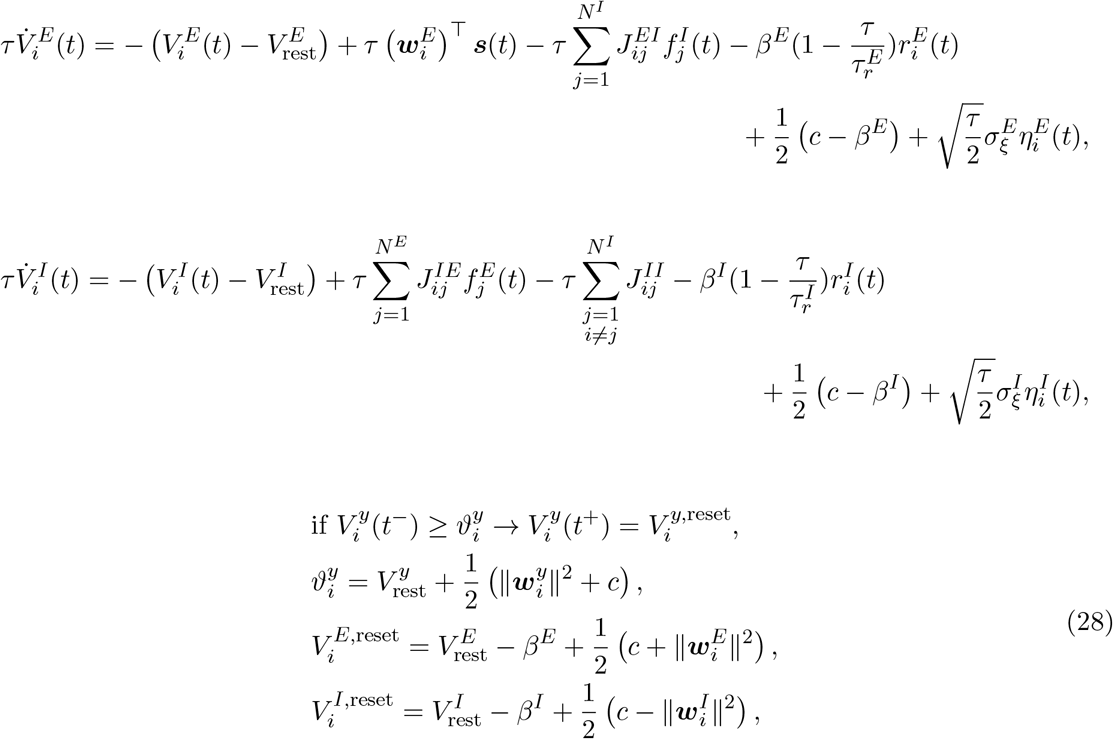

where 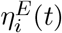 and 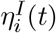 are the independent Gaussian white noise processes defined in Eq. (13) above. We note that all terms on the right-hand side of Eq. (28) have the desired units of mV. The model in Eq. (28) is an efficient E-I spiking network with improved compatibility with neurobiology. We have expressed two new terms in the membrane potentials of E and I neurons, one dependent on the metabolic constant *β*^*y*^ and one on the noise that we assumed in the condition for spiking (see Eq. 12). We will group these two terms to define an external current, a current that is well known in spiking models of neural dynamics ^41^.

### Efficient generalized leaky integrate-and-fire neuron model

Finally, we rewrite the model from Eq. (28) in a compact form in terms of transmembrane currents, and discuss their biological interpretation. The efficient coding with spikes is realized by the following model for the neuron *i* of type *y* ∈ {*E, I*}:

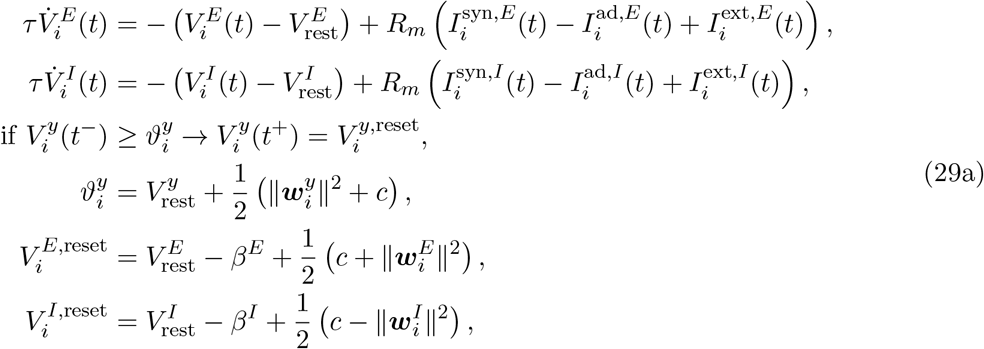

with *R*_*m*_ the current resistance. The leak current,

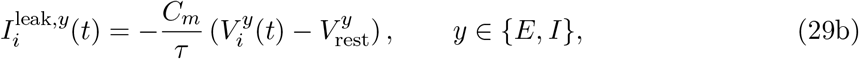

with *τ* = *R*_*m*_*C*_*m*_ and *C*_*m*_ the capacitance of the neural membrane^64^, arose by assuming the same time constant for the target signals ***x***(*t*) and estimates 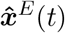 and 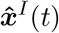 (see Eqs. 8 and 18). We see that the passive membrane time constant *τ* = λ^−1^ can be traced back to the time constant of the population read-out in Eq. (9). The synaptic currents are defined as

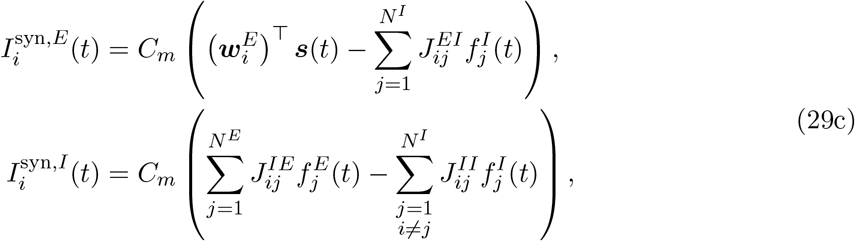

where we note the presence of a feedforward current to E neurons,

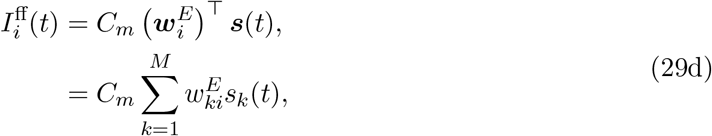

which consist in a linear combination of the stimulus features ***s***(*t*) weighted by the decoding weights 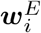. The stimulus features can be traced back to the definition of the target signals in Eq. (8). This current emerges in E neurons, as a consequence of having the target signal ***x***(*t*) in the loss function of the E population (see Eqs. 10-11). I neurons do not receive the feedforward current because their loss function does not contain the target signal.

The current providing within-neuron feedback triggered by each spike,

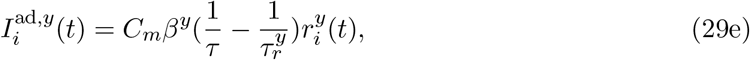

was recently recovered ^38^. This current has the kinetics of the single neuron readout 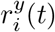 (i.e., low-pass filtered spike train). Its sign depends on the relation between the time constant of the population readout *τ* = λ^−1^ and single neuron readout 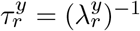, because the metabolic constant *β*^*y*^ is non-negative by definition (Eq. 10). If the single neuron readout is slower than the population readout, 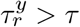, within-neuron feedback is negative, and can thus be interpreted as spike-triggered *adaptation*. On the contrary, if the single neuron readout is faster than the population readout, 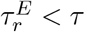, the within-neuron feedback is positive and can thus be interpreted as spike-triggered *facilitation*. In a special case where the time constant of the single neuron and population readout are assumed to be equal, within-neuron feedback vanishes.

Finally, we here derived the non-specific external current:

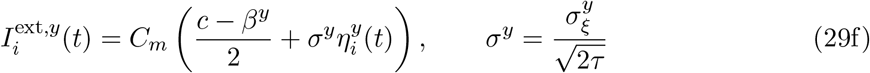

that captures the ensemble of non-specific synaptic currents received by each single neuron. The non-specific current has a homogeneous mean across all neurons of the same cell type, and a neuron-specific fluctuation. The mean of the non-specific current can be traced back to the weighting of the metabolic cost over the encoding error in model objectives (Eq. 10), while the fluctuation can be traced back to the noise strength that we assumed in the condition for spiking (Eq. 12). The non-specific external current might arise because of synaptic inputs from other brain areas than the brain area that delivers feedforward projections to the E-I network we consider here, or it might result from synaptic activity of neurons that are part of the local network, but are not tuned to the feedforward input ^86^.

We also recall the fast and slower time scales of single neuron activity:

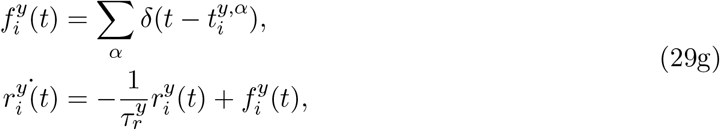

and the connectivity matrices

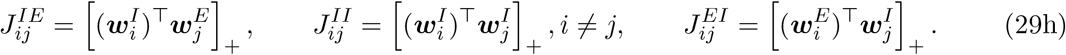

The structure of synaptic connectivity is fully determined by the similarity of tuning vectors of the presynaptic and the postsynaptic neurons (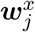 and 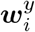) respectively, while the distribution of synaptic connectivity weights is fully determined by the distribution of tuning parameters 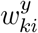.

### Stimulus features

We define stimulus features ***s***(*t*) as a set of *k* = 1, …, *M* independent Ornstein-Uhlenbeck processes with vanishing mean, standard deviation *σ*^*s*^ and the correlation time 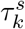,

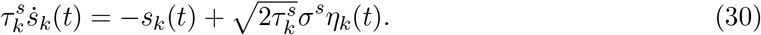

If not mentioned otherwise, we use the following parameters, identical across stimulus features: *σ*^*s*^ = 2 (mV)^1/2^ and 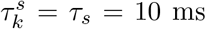. Variables *η*_*k*_(*t*) are independent Gaussian white noise processes with zero mean and covariance function ⟨*η*_*k*_(*t*)*η*_*l*_(*t*^*′*^)⟩=*δ*_*kl*_*δ*(*t* − *t*^*′*^). These variables should not be confused with the Gaussian white noises 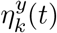 in Eq. (28).

### Parametrization of synaptic connectivity

In the efficient E-I model, synaptic weights 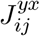 are parametrized by tuning parameters 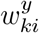 through Eq. (24). The total number of synapses in the E-I, I-I and I-E connectivity matrices (including silent synapses with zero synaptic weight) is *n*_syn_ = 2*N* ^*E*^*N* ^*I*^ + (*N* ^*I*^)^2^, while the number of tuning parameters is *n*_*w*_ = *M* (*N* ^*E*^ + *N* ^*I*^). Because the number of stimulus features *M* is expected to be much smaller than the number of E or I neurons, the number of tuning parameters *n*_*w*_ is much smaller than the number of synapses *n*_syn_.

We can achieve a further substantial decrease in the number of free parameters by using a parametric distribution of tuning parameters 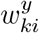. We set the tuning parameters following a normal distribution and found that excellent performance can be achieved with random draws of tuning parameters from the normal distribution, thus without searching for a specific set of tuning parameters. This drastically decreased the number of free parameters relative to synaptic weights to only a handful of parameters that determine the distributions of tuning parameters.

Given *M* features, we sampled tuning parameters, 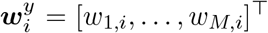, with *i* = 1, …, *N* ^*y*^, *y* ∈ {*E, I*}, as random points uniformly distributed on a *M*-dimensional sphere of radius 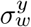. We obtained this by sampling, for each neuron, a vector of *M* i.i.d. standard Gaussian random variables, 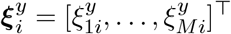, with 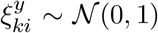, and normalizing the vector such as to have length equal to 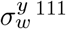,

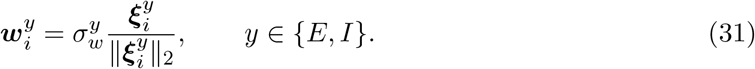

This ensures that the length of tuning vectors 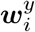 in Eq. (31) is homogeneous across neurons of the same cell type, i.e., 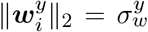. Parameters 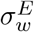 and 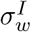 determine the heterogeneity (spread) of tuning parameters.

By combining Eq. (24) and Eq. (31), we obtain the synaptic weights, 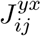, as a function of the angle, 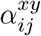, between the tuning vectors of presynaptic neurons, 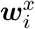, and postsynaptic neurons, 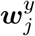,

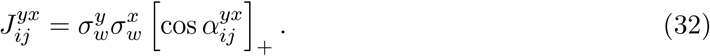

In the *M* = 3 dimensional case, we have that the distribution of the angle between two vectors is 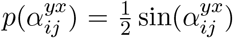, with 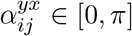. Thus, the average strength of synaptic weights between the pre- and the postsynaptic population can be calculated as

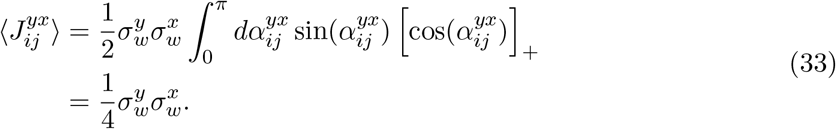

Thus, the upper bound for the synaptic weight between cell types *x* and *y* is simply

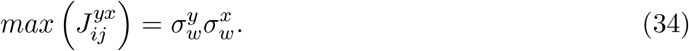

From the Eq. (33), we have that the mean E-I connectivity is equal to the mean I-E connectivity, 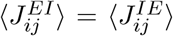. As we consider the ratio of the mean connectivity between I-I and E-I connections, we find that it is given by the following:

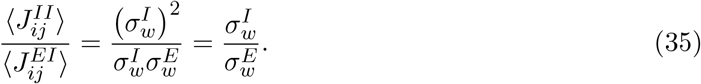

## Performance measures

### Average encoding error and average metabolic cost

The definition of the time-dependent loss functions (Eq. 10) induces a natural choice for the performance measure: the mean squared error (MSE) between the targets and their estimators for each cell type. In the case of the E population, the time-dependent encoding error is captured by the variable *ϵ*^*E*^(*t*) in the Eq. (11) and in case of I population it is captured by *ϵ*^*I*^(*t*) defined in the same equation. We used the root MSE (RMSE), a standard measure for the performance of an estimator ^41^. For the cell type *y* ∈ {*E, I*} in trial *q*, the RMSE is measured as

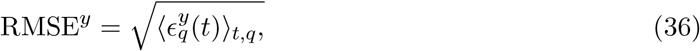

with ⟨*z*_*q*_(*t*)⟩_*t,q*_ denoting the time- and trial-average.

Following the definition of the time-dependent metabolic cost in the loss functions (Eq. 10), we measured the average metabolic cost in a trial *q* for the cell type *y* ∈ {*E, I*} as

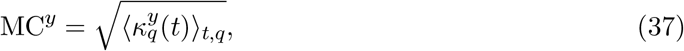

with time-dependent metabolic cost *κ*^*y*^(*t*) as in model’s objectives (Eq. 11) and ⟨*z*_*q*_(*t*)⟩_*t,q*_ the time- and trial-average. The square root was taken to have the same scale as for the RMSE (see Eq. 36).

### The bias of the estimator

The MSE can be decomposed into the bias and the variance of the estimator. The time-dependent bias of estimates 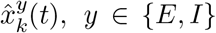, were evaluated for each time point over *q* = 1, …, *Q* trials. The time-dependent bias in input dimension *k* = 1, …, *M* is defined as

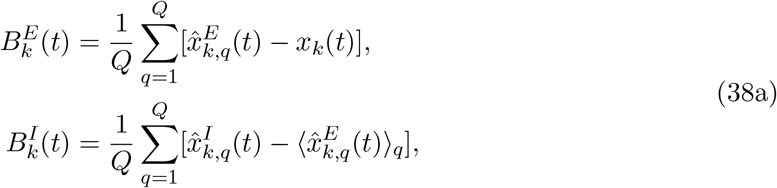

with ⟨*z*_*q*_(*t*)⟩_*q*_ the trial-averaged realization at time *t*. To have an average measure of the encoding bias, we averaged the bias of estimators over time and over input dimensions:

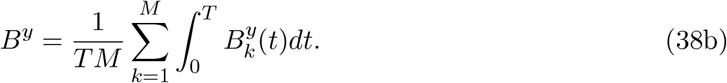

The averaging over time and input dimensions is justified because *s*_*k*_(*t*) are independent realizations of the Ornstein-Uhlenbeck process (see Eq.30) with vanishing mean and with the same time constant, and variance across input dimensions.

### Criterion for determining optimal model parameters

The equations of the E-I spiking network in Eqs. 29a-29h (Methods), derived from the instantaneous loss functions, give efficient coding solutions valid for any set of parameter values. However, to choose parameters values in simulated data in a principled way, we performed a numerical optimization of the performance function detailed below. Numerical optimization gave the set of optimal parameters listed in Table 1. When testing the efficient E-I model with simulations, we used the optimal parameters in Table 1 and changed only the parameters plotted in the figure axes on a figure-by-figure basis.

To estimate the optimal set of parameters *θ* = *θ*^*^, we performed a grid search on each parameter *θ*_*i*_ while keeping all other parameters fixed as specified in Table 1. While varying the parameters, we measured a weighted sum of the time- and trial-averaged encoding error and metabolic cost. For each cell type *y* ∈ {*E, I*}, we computed

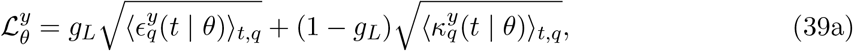

with ⟨*z*_*q*_(*t*)⟩_*t,q*_ the average over time and over trials and with *ϵ*^*y*^(*t*) and *κ*^*y*^(*t*) as in model’s objectives (Eq. 11), where *g*_*L*_ ∈ [0, 1] is a weighting factor.

To optimize the performance measure, we used a value of *g*_*L*_ = 0.7. The parameter *g*_*L*_ in the Eq. (39a) regulates the relative importance of the average encoding error over the average metabolic cost. Since the performance measure in Eq. (39a) is closely related to the average over time and trials of the instantaneous loss function (Eq. 10) where the parameter *β* regulates the relative weight of instantaneous encoding error over the metabolic cost, setting *g*_*L*_ is effectively achieved by setting *β*.

The optimal parameter set *θ* = *θ*^*^ reported in Table 1 is the parameter set that minimizes the sum of losses across E and I cell type

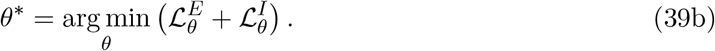

For visualization of the behavior of the average metabolic cost (Eq. 37) and average loss (Eq. 39a) across a range of a specific parameter *θ*_*i*_, we summed these measures across the E and I cell type and normalized them across the range of tested parameters.

The exact neural dynamics and performance of our model depends on the realizations of random variables which describe the the tuning parameters 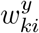, the Gaussian noise in the non-specific currents 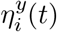, and the initial conditions of the membrane potential 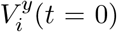, that were randomly drawn from a normal distribution in each simulation trial. To capture the performance of a “typical” network, we iterated the performance measures across trials with different realizations of these random variables, and averaged the performance measures across trials. We typically used 100 simulation trials for each parameter value.

## Functional activity measures

### Tuning similarity

The pairwise tuning similarity was measured as the cosine similarity^112^, defined as:

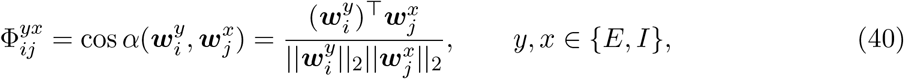

with 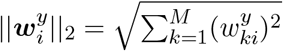 the length of the tuning vector in Euclidean space and *α* the angle between the tuning vectors 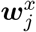 and 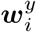.

### Cross-correlograms of spike timing

The time-dependent coordination of spike timing was measured with the cross-correlogram (CCG) of spike trains, corrected for stimulus-driven coincident spiking. The raw cross-correlogram (CCG) for neuron *i* of cell type *y* and neuron *j* of cell type *x* was measured as follows:

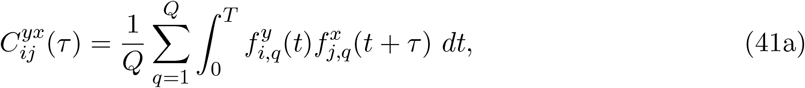

with *q* = 1, …, *Q* simulation trials with identical stimulus and *T* the duration of the trial. We subtracted from the raw CCG the CCG of trial-invariant activity. To evaluate the trial-invariant cross-correlogram, we first computed the peri-stimulus time histogram (PSTH) for each neuron as follows:

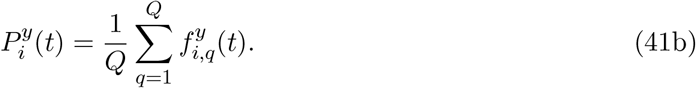

The trial-invariant CCG was then evaluated as the cross-correlation function of PSTHs between neurons *i* and *j*,

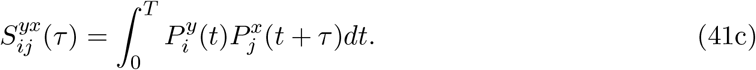

Finally, the temporal coordination of spike timing was computed by subtracting the correction term from the raw CCG:

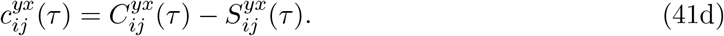

### Average imbalance of synaptic inputs

We considered time and trial-averaged synaptic inputs to each E and I neuron *i* in trial *q*, evaluated as:

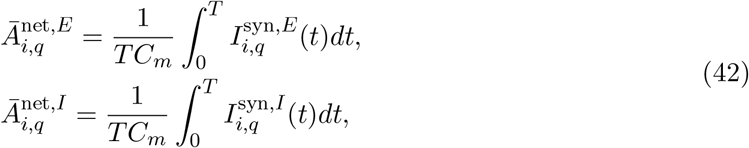

with synaptic currents to E neurons 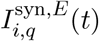 and to I neurons 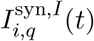 as in Eq. (29c). Synaptic inputs were measured in units of mV. We reported trial-averages of the net synaptic inputs from the Eq. (42).

### Instantaneous balance of synaptic inputs

We measured the instantaneous balance of synaptic inputs as the Pearson correlation of time-dependent synaptic inputs incoming to the neuron *i*. For those synaptic inputs that are defined as weighted delta-spikes (for which the Pearson correlation is not well defined; see Eq. 29c), we convolved spikes with a synaptic filter 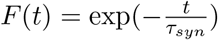,

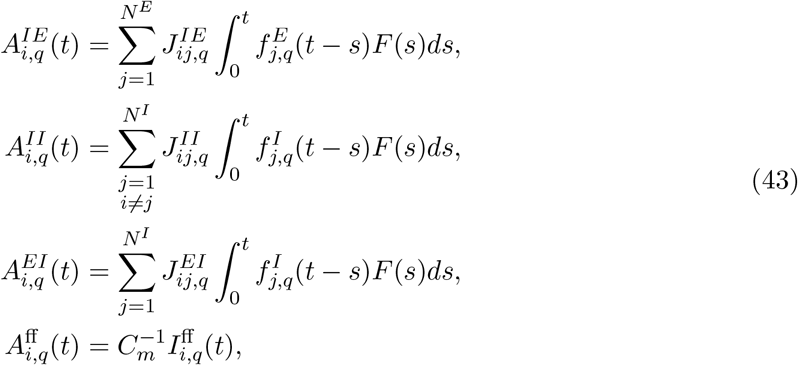

where we used the expression for the feedforward synaptic current from the Eq. (29d). Note that the feedforward synaptic current is already already low-pass filtered (see Eq. 30). Using synaptic inputs from the Eq. 43, we computed the Pearson correlation of synaptic inputs incoming to single E neurons, 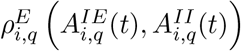 for *i* = 1, …, *N* ^*E*^, and to single I neurons, 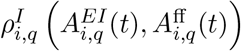 for *i* = 1, …, *N* ^*I*^. The coefficients were then averaged across trials.

### Tuning curves and selectivity index

The selectivity index of a neuron captures the change in neuron’s firing rate in response to a change in the stimulus. We first evaluated the tuning curve of each neuron by measuring the firing rate of the neuron 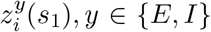, as a function of the amplitude of the stimulus feature *s*_1_. The firing rate was evaluated from the network response to *M* = 3 stimulus features that were constant over time. We varied the first stimulus feature *s*_1_ from strongly negative (*s*_1_ = −5) to strongly positive values (*s*_1_ = *s*_max_ = 5), while the two other features were kept at an intermediate positive value (*s*_2_ = *s*_3_ = 1.6). Note that with all three features at such intermediate value (*s*_1_ = *s*_2_ = *s*_3_ = 1.6), the average firing rate was about 8 Hz in E and 12 Hz in I neurons. To evaluate the tuning curve of a neuron, we measured its firing rate in 100 simulation trials of 1 second duration, for each value of the stimulus feature *s*_1_.

To evaluate the sensitivity index, we normalized the tuning curve of the neuron with its maximal value,

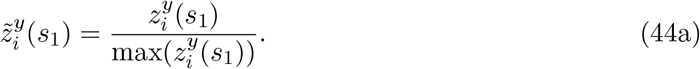

We then computed the sensitivity index as the average absolute change of the normalized firing rate with the change in the stimulus:

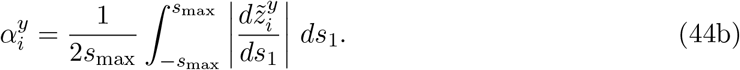

## Perturbation experiments

### Perturbation of neural activity

Empirical studies ^58,59^ suggested experiments with perturbation of neural activity that estimate functional connectivity in recurrently connected neural networks. Here, we detail the procedure on how we performed similar experiments on simulated neural networks. To evaluate the functional connectivity between pairs of neurons, we measured the effect of activation of a single E neuron (“target” neuron) on the activity of other neurons. We stimulated a randomly chosen E neuron with a depolarizing input, capturing the effect of photostimulation in empirical studies ^58,59^, and measured the deviation of the firing rate from the baseline in all other neurons.

The time-dependent deviation of the firing rate from the baseline for neuron *i* of type *y* ∈ {*E, I*} was computed as 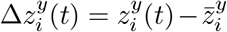, with 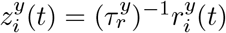 the estimate of the instantaneous firing rate and 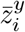 the average spontaneous firing rate of the neuron *i*. The target neuron received a constant depolarizing current during 50 ms and the effect of its activity on other neurons was measured during a time window of [0, 100] ms with respect to the onset of the stimulation. The functional connectivity between the target neuron and every other neuron in the network was then computed as the time average of the variable 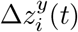. To isolate the functional effect of recurrent connections on firing rate changes, we performed these experiments in a network without external stimuli, setting *s*_*k*_(*t*) = 0 ∀*t, k*.

#### Removal of connectivity structure

To better understand the effect of optimally structured recurrent connectivity (as given by the Eq. 24) on network’s activity and efficiency, we compared networks with and without the connectivity structure. To fully remove the connectivity structure, we randomly permuted, without repetition, recurrent connectivity weights between all neuronal pairs of all the three recurrent connectivity matrices. This was achieved by shuffling entries within each recurrent connectivity matrix. This procedure preserves all properties of the distribution of connectivity weights and only removes the connectivity structure. Shuffling of connections was iterated across 200 simulation trials, with each trial implementing a different random permutation of the connectivity. Dale’s law is preserved by such manipulation.

To compare the performance of models with structured and unstructured connectivity (as reported on Fig. 4A), we collected the low-pass filtered spiking activity in networks with and without connectivity structure. We used this neural activity to train a linear decoder with least squares method that minimizes the Euclidean distance between target signals and a linear readout of low-pass filtered spikes. The output of the training was a set of linear coefficients akin to decoding weights 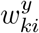. We used these decoding weights estimated by the decoder to weight spikes in a held-out validation set. The performance was measured with root mean squared error (RMSE) between target signals and their estimates in the validation set. The training set comprised 70 % of trials (140 trials), and the validation test comprised the remaining 30 % of trials (60 trials).

To compare networks with and without connectivity structure about their metabolic cost, firing rate, variability of spiking and the E-I balance (Fig. 4B-G), we performed these measures in networks with and without connectivity structure and plotted their distributions across 200 simulation trials. For the comparison of the metabolic cost (Fig. 4B), we additionally matched the network with and without the connectivity structure about their mean net synaptic input to E and I neurons, to see if the difference in the metabolic cost between structured and unstructured networks persists after such matching. For the comparison of the coefficient of variation in structured and unstructured networks (Fig. 4E), we used a constant stimulus instead of the OU stimulus, to exclude possible effects of a time-dependent variations of the stimulus on the variability of spiking. Constant stimulus was homogeneous across all stimulus dimensions, *s*_*k*_(*t*) = 1.6, ∀*k* = 1, …, *M*. The amplitude of the constant stimulus was set such that the average firing rate in response to the constant stimulus matched the firing rate in response to the OU stimulus.

For the comparison of the voltage correlations and the effective connectivity between structured and unstructured networks (Fig. 4H-I), we additionally permuted individual connectivity (sub)matrices. This gave four cases, namely, permuted E-I, I-I, I-E, and “all”, with “all” meaning that all three recurrent connectivity matrices have been randomly permuted.

We also tested networks where the connectivity structure was not fully but only partially removed. There, we limited random permutation of synaptic weights to pairs of neurons that already had a connection in the structured network. By the Eq. 24, connected neurons are those with positive tuning similarity, i.e., neuronal pairs for which the following holds: 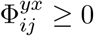, with tuning similarity as in Eq. 40. We compared partially unstructured networks with structured networks by plotting measures of neural activity in structured and partially unstructured networks across 200 simulation trials (Fig. S3B-E).

#### Perturbation of connectivity

To test the robustness of the model to random perturbations of synaptic weights (Fig. S3G-H), we applied a random jitter to optimally efficient recurrent synaptic connectivity weights. The random jitter was proportional to the optimal synaptic weight, 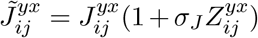, where *σ*_*J*_ is the strength of the perturbation and 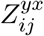 are independent standard normal random variables. All three recurrent connectivity matrices (E-I, I-I and I-E) were randomly perturbed at once.

#### Computer simulations

We ran computer simulations with Matlab R2023b (Mathworks). The membrane equation for each neuron was integrated with Euler integration scheme with the time step of *dt* = 0.02 ms.

The simulation of the E-I network with 400 E units and 100 I units for an equivalent of 1 second of neural activity lasted approximately 1.65 seconds on a laptop.

## Supplementary material

### Supplementary text 1: Derivation of the one cell type model

An efficient spiking model network with one cell type (1CT) has been developed previously ^28^, and properties of the 1CT model where the computation is assumed to be the leaky integration of inputs has been addressed in a number of previous studies ^29,37,36,33,43^. Compared to the efficient E-I model, the 1CT model can be seen as a simplification, and can be treated similarly to the E-I model, which is what we demonstrate in this section.

As the name of the model suggests, all neurons in the 1CT model are of the same cell type, and we have *i* = 1, …, *N* such neurons. We can then use the definitions in Eqs. (6) – (9) (now without the index *y*) and a loss function similar to the one in ^36^, but with only one (quadratic) regularizer

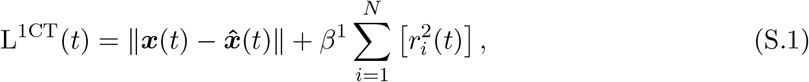

with *β*^1^ *>* 0. The encoding error of the 1CT model minimizes the squared distance between the target signal *x*(*t*) and the estimate 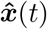. As we apply the condition for spiking as for the E-I network (Eq. 12 without the index *y*) and follow the same steps as for the E-I network, we get

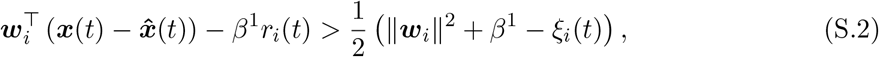

with *ξ*_*i*_(*t*) the noise at the condition for spiking. Same as in the E-I model, we define the noise as an Ornstein-Uhlenbeck process with zero mean, obeying

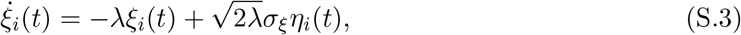

where *η*_*i*_ is a Gaussian white noise and λ = *τ*^−1^ is the inverse time constant of the process. We now define proxies of the membrane potential and the firing threshold as

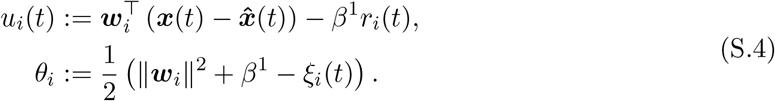

Differentiating the proxy of the membrane potential *u*_*i*_(*t*) and rewriting the model as an integrate-and-fire neuron, we get

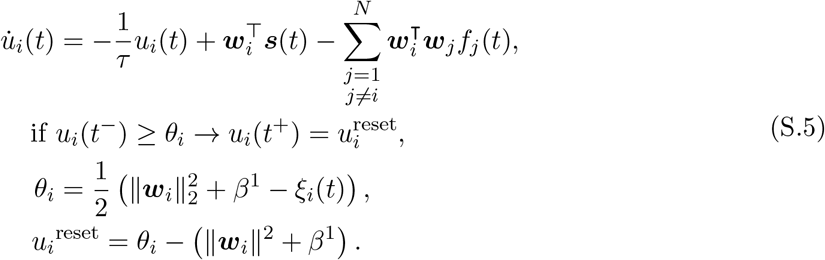

We now proceed in the same way as with the E-I model and define new variables

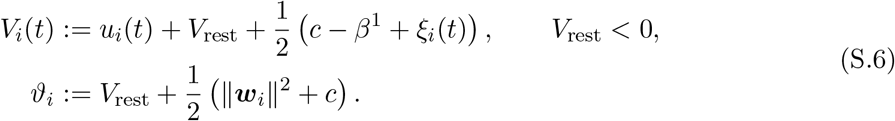

In these new variables, we can rewrite the membrane equation of the 1CT model as follows:

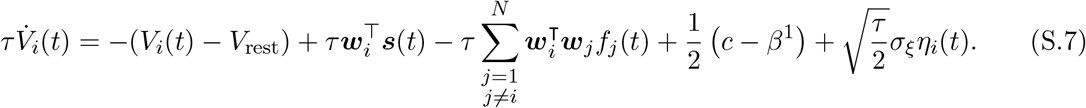

Finally, we rewrite the model with a more compact notation of a leaky integrate-and-fire neuron model with transmembrane currents,

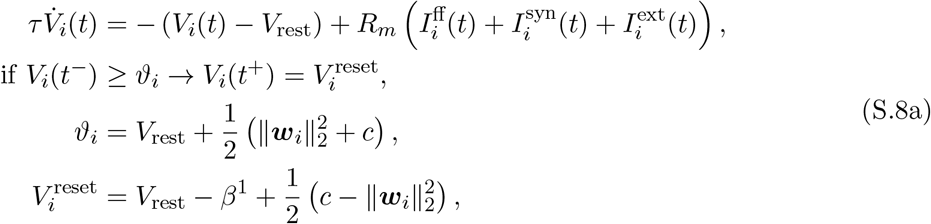

with currents

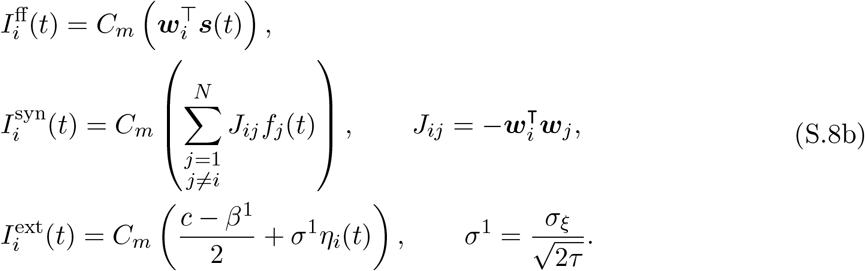

Note that the model with one cell type does not obey Dale’s law, since the same neuron sends to its postsynaptic targets excitatory and inhibitory currents, depending on the tuning similarity of the presynaptic and the postsynaptic neuron ***w***_*i*_ and ***w***_*j*_ (Eq. S.8b). In particular, if the pre- and postsynaptic neurons have similar selectivity 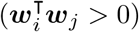, the recurrent interaction is inhibitory, and if the neurons have different selectivity 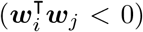, the interaction is excitatory. Simply put, neurons with similar selectivity inhibit each other while neurons with different selectivity excite each other ^36^.

Dale’s law can be imposed to the 1CT model the same way as in the E-I model, by removing synaptic interactions between neurons with different selectivity with rectification of the connectivity matrix,

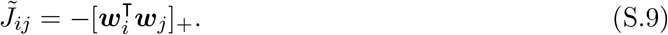

However, this manipulation results in a network with only inhibitory recurrent synaptic interactions, and thus a network of only inhibitory neurons. Network with only inhibitory interactions is less relevant for the description of recurrently connected biological networks.

### Supplementary text 2: Parameters of the E-I model without non-specific currents

**Table S1.** Table of optimal model parameters for the efficient E-I network without non-specific synaptic currents. As in Table 1, for the E-I model without non-specific currents. The model is defined in Eq. 25.

| parameter | notation | value |
| --- | --- | --- |
| number of E neurons | $N^E$ | 400 |
| ratio of E to I neuron numbers | $N^E : N^I$ | 4:1 |
| number of the input features | $M$ | 3 |
| time constant of the population readout (E and I) | $\tau$ | 10 ms |
| time constant of the single neuron readout | $\tau_r^E = \tau_r^I$ | 10 ms |
| noise strength (non-specific current) | $\sigma$ | 5.0 mV |
| heterogeneity factor of tuning parameters in E | $\sigma_w^E$ | $1.0 \text{ (mV)}^{\frac{1}{2}}$ |
| ratio of mean E-I to I-I synaptic connectivity | mean E-I : mean I-I | 3:1 |
| metabolic constant | $\beta$ | 14 mV |
| (threshold constant) | $(c/2)$ | (0 mV) |
| distance threshold to reset potential (E neurons) | $\vert \vartheta^E - V_{\text{rest}}^E\vert $ | 7.5 mV |
| distance threshold to reset potential (I neurons) | $\vert \vartheta^I - V_{\text{rest}}^I\vert $ | 11.5 mV |
| connection probability (recurrent synapses) | $p^{IE} = p^{II} = p^{EI}$ | 0.5 |
| mean E-I synaptic weight (EPSP to I at max) | $\langle J_{ij}^{IE} \rangle$ | 0.75 mV |
| mean I-E synaptic weight (IPSP to E at max) | $\langle J_{ij}^{EI} \rangle$ | 0.75 mV |
| mean I-I synaptic weight (IPSP at max) | $\langle J_{ij}^{II} \rangle$ | 2.25 mV |

Our analytical derivation in Eq. 25 suggested an efficient E-I model that is simpler with respect to the E-I model studied in this contribution, as it does not have non-specific synaptic currents. Optimal (computational) model parameters of such simpler model, listed above the double line in Table S1, are by definition identical to the full E-I model listed in Table 1. However, the model without non-specific synaptic currents differs from the full E-I model about the distance between the resting potential and the threshold. In the simpler model, this distance is lower compared to the full E-I model, and is not consistent with empirically measured distance, which is about 20 mV ^63^.

A simple way to increase the distance between the resting potential and the firing threshold is to introduce a constant that multiplies all mathematical terms in the Eq. 25. While this allows to achieve biologically plausible values for the distance between the resting potential and the threshold, it leads to values of mean recurrent synaptic connectivity 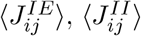 and 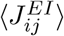 that are stronger than typically reported in the empirical literature ^61^.

### Supplementary text 3: Analysis of the one cell type model and comparison with the E-I model

We re-derived the 1CT model as a simplification of the E-I network (Supplementary Text 1, Supplementary Fig. S1A-B), with objective function of the same form as *L*^*E*^ and by allowing a single type of neurons sending both excitatory and inhibitory synaptic currents to their post-synaptic targets (Supplementary Fig. S1C). Similarly to the E-I model, also the 1CT model exhibits structured connectivity, with synaptic strength depending on the tuning similarity between the presynaptic and the postsynaptic neuron. Pairs of neurons with stronger tuning similarity (dissimilarity) have stronger mutual inhibition (excitation); see Supplementary Fig. S1D.

We compared the coding performance of the E-I model with that of a fully connected 1CT model. Both models received the same set of stimulus features and performed the same computation. In the 1CT model, tuning parameters were drawn from the same distribution as used for the E neurons in the E-I model. We used the same membrane time constant *τ* in both models, while the metabolic constants (*β* of the E-I model and *β*^1^ of the 1CT model) and the noise strength (*σ* of the E-I model and *σ*^1^ of the 1CT model) were chosen such as to optimize the average loss for each model (Fig. 6B for E-I model, Supplementary Fig. S1F-G for 1CT model). Parameters of the 1CT model are listed in the Supplementary Table S2. A qualitative comparison of the E-I and the 1CT model showed that with optimal parameters, both models accurately tracked multiple target signals (Fig. 1G and Supplementary Fig. S1E).

To compare the performance of the E-I and the 1CT models also quantitatively, we measured the average encoding error (RMSE), metabolic cost (MC) and loss of each model. The RMSE and the MC in the 1CT model were measured as in Eq. 36 and 37, while the average loss of each model was evaluated as follows:

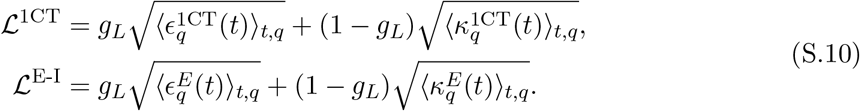

Unless mentioned otherwise, we weighted stronger the encoding error compared to the metabolic cost and used *g*_*L*_ = 0.7.

**Table S2.** Table of default model parameters for the efficient network with one cell type. The parameters *N,M,τ* and 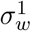 were chosen identical to the E-I network (see Table 1 in the main text). Parameters *σ*^1^ *β*^1^ were determined as values that maximize network efficiency (see section “Performance measures” in the main text).

| parameter | notation | value |
| --- | --- | --- |
| number of E neurons | $N$ | 400 |
| number of the input features | $M$ | 3 |
| time constant of the single neuron and population readout | $\tau$ | 10 ms |
| noise strength | $\sigma^1$ | 1.8 mV |
| SD of tuning parameters | $\sigma_w^1$ | 1 (mV)<sup>1/2</sup> |
| metabolic constant | $\beta^1$ | 11.4 mV |

## Supplementary Figures

**Figure S1.**
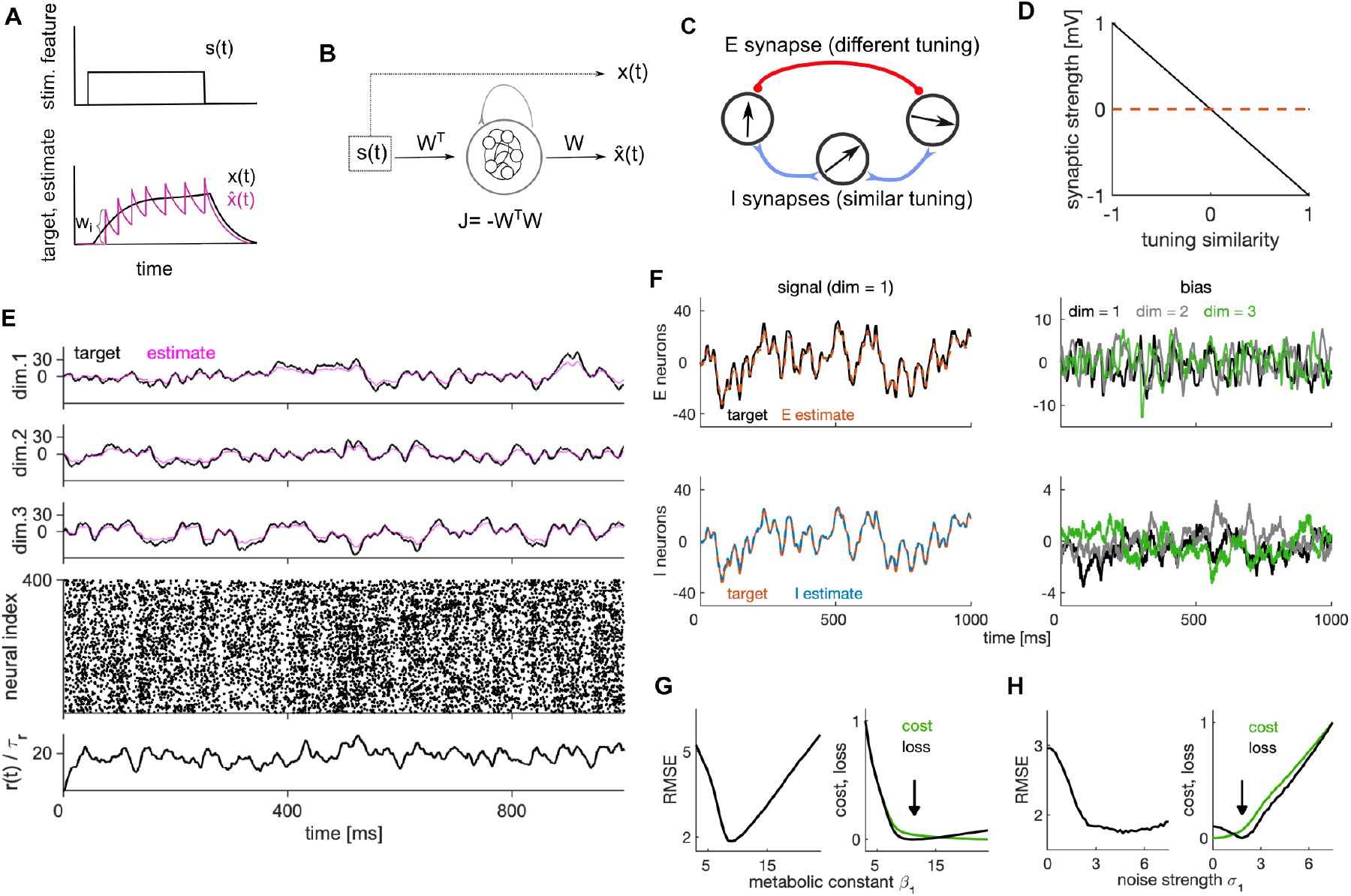
Efficient spiking model with one cell type and the encoding bias of the E-I network. **(A)** Schematic of efficient coding with a single spiking neuron. In this toy example, the neuron has a positive decoding weight and responds to a single stimulus feature *s*(*t*) (top). The target signal *x*(*t*) (bottom, black) integrates the stimulus feature from top. The neuron spikes to keep the readout of its activity 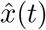(magenta) close to the target. **(B)** Schematic of an efficient 1CT model. The target signal is computed from the stimulus feature. The spiking network estimates the targets by generating population readouts of its spiking activity. **(C)** Schematic of excitatory (red) and inhibitory (blue) synaptic interactions in the 1CT model. Neurons with similar selectivity inhibit each other (blue), while neurons with different selectivity excite each other (red). The same neuron is sending excitatory and inhibitory synaptic outputs, which is not consistent with Dale’s law. **(D)** Strength of recurrent synapses as a function of pair-wise tuning similarity. **(E)** Simulation of the model with 1CT. Top three rows show the target (black), and the estimate (magenta) in each of the 3 input dimensions. **(F)** Top left: The target (black) and the estimate (red) of the E population in the first signal dimension (in response to the first stimulus feature *s*_1_(*t*)). The estimate is averaged across 100 trials, with trials varying about the initial conditions and the noise in the membrane potential. Bottom left: Same as on top left, for the I population. Right: Time-dependent bias of estimates in E (top) and I (bottom) population in each of the three stimulus dimensions. **(G)** Left: Root mean squared error (RMSE) as a function of the metabolic constant *β*^1^. Right: Normalized metabolic cost (green) and normalized average loss (black) as a function of the metabolic constant *β*^1^. The black arrow denotes the minimum of the loss and thus the optimal parameter *β*^1^. **(H)** Same as in **G**, measured as a function of the noise strength *σ*^1^. Results in F,G and H were computed in 100 simulation trials of duration of 1 second. For other parameters, see Table 1 (E-I model) and Table S2 (1CT model). This figure is related to Fig. 1 in the main paper.

**Figure S2.**
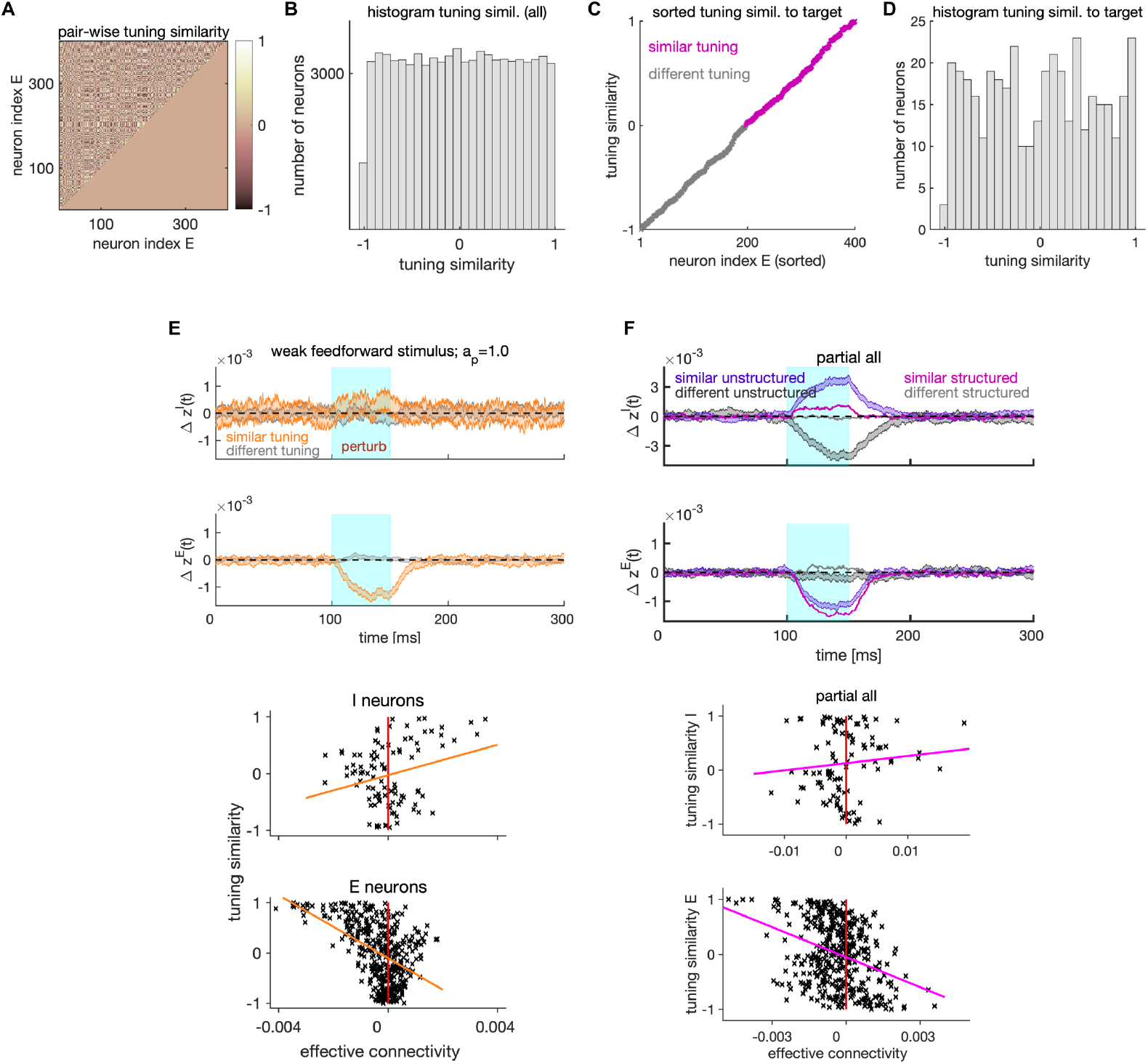
Tuning similarity and its relation to lateral excitation/inhibition. **(A)** Pair-wise tuning similarity for all pairs of E neurons. Tuning similarity is measured as cosine similarity of decoding vectors between the target neuron and every other E neuron. **(B)** Histogram of tuning similarity across all E-E pairs shown in **A.(C)** Tuning similarity to a single, randomly selected target neuron. Tuning similarity to a target neuron corresponds to a vector from the tuning similarity matrix in **A**. We sorted the tuning similarity to target from the smallest to the biggest value. Neurons with negative similarity are grouped as neurons with different tuning, while neurons with positive tuning similarity are grouped as neurons with similar tuning. **(D)** Histogram of tuning similarity of E neurons to the target neuron shown in **C**. With distribution of tuning parameters that is symmetric around zero as used in our study, any choice of the target neuron gives approximately the same number of neurons with similar and different selectivity. **(E)** Top: Trial and neuron-averaged deviation of the instantaneous firing rate from the baseline in presence of weak feedforward stimulus. We show the ± standard error of the mean (SEM) of neurons with similar (orange) and different tuning (gray) to the target neuron. The photostimulation intensity is at threshold (*a*_*p*_ = 1.0). The feedforward stimulus was received by all E neurons and it induced, together with the external current, the mean firing rates of 7.3 Hz and 13.5 Hz in E and I neurons, respectively. Bottom: Scatter plot of the tuning similarity versus effective connectivity. Magenta line marks the least-squares line. **(F)** Same as in **E**, for the network with partial (fine-grained) removal of connectivity structure. Partial removal of connectivity structure is achieved by shuffling the synaptic weights among pairs of neurons with similar tuning 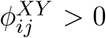. For model parameters, see Table 1. This figure is related to Fig. 3 in the main paper.

**Figure S3.**
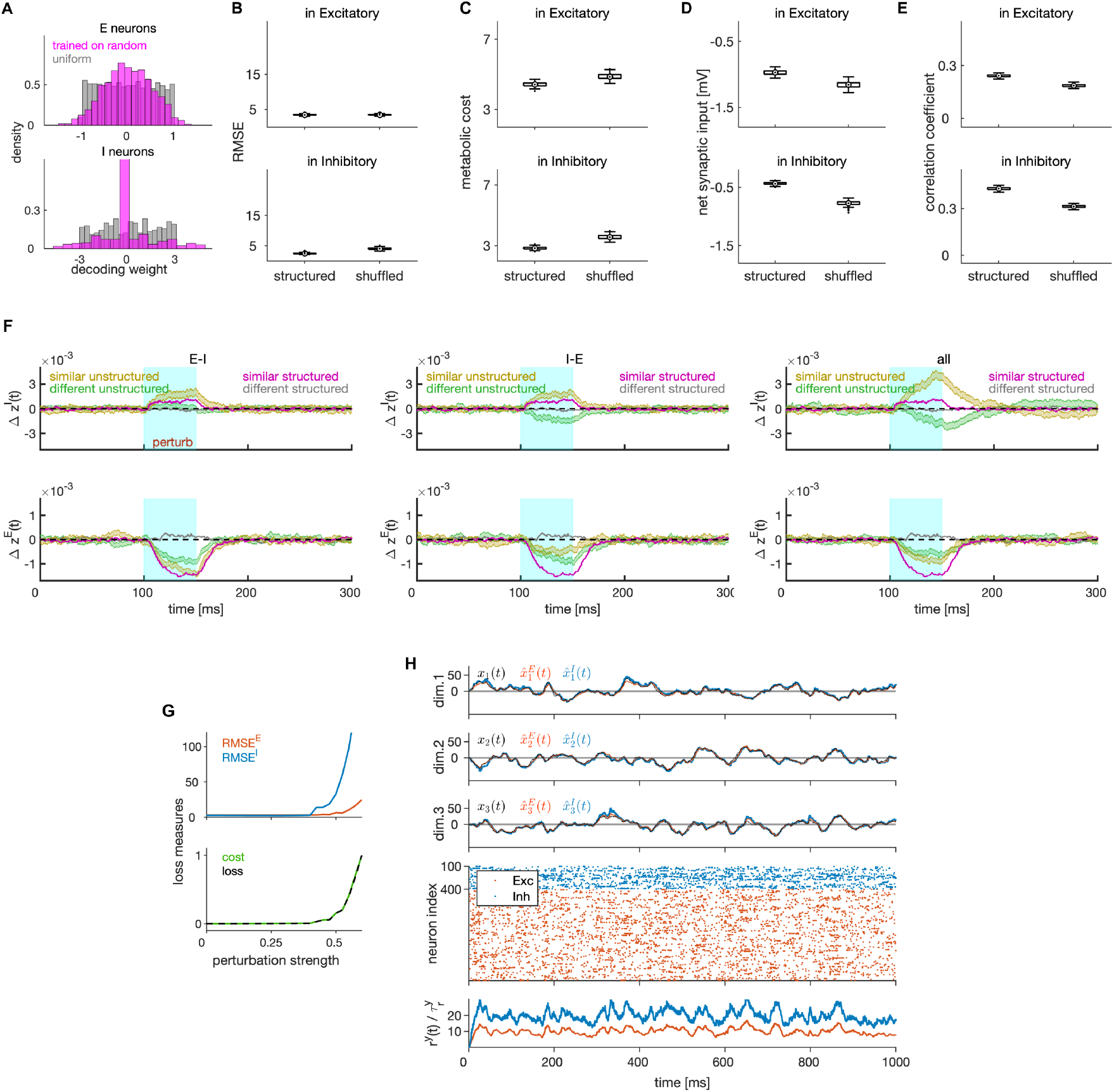
Effect of removal of connectivity structure and of jittering of synaptic weights. **(A)** Distribution of decoding weights after training a linear decoder on neural activity generated by the network without connectivity structure. **(B)** RMSE in E (top) and I neurons (bottom) in networks with partial removal of connectivity structure. Partial removal of connectivity structure is achieved by limiting the permutation of synaptic connectivity to neuronal pairs with similar tuning, e.g. to neuronal pairs for which the following is true: 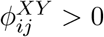. **(C)** Same as in **B**, showing the average metabolic cost on spiking. **(D)** Same as in **B**, showing the average net synaptic input, a measure of the average E-I balance. **(E)** Same as in **B**, showing the correlation of synaptic inputs, a measure of instantaneous balance. **(F)** Average deviation of the instantaneous firing rate from the baseline for the population of I (top) and E (bottom) neurons in networks with fully removed structure in E-I (left), I-E (middle) and in all connectivity matrices (right). We show the mean ± SEM for neurons with similar (ochre) and different (green) tuning to the target neuron. The mean traces of the network with structured connectivity are shown for comparison, with magenta and gray for similar and different tuning, respectively. **(G)** Top: The RMSE (top) in E and I cell type, as a function of the strength of perturbation of the synaptic connectivity by random jittering. Bottom: Same as on top, showing the normalized metabolic cost (green) and average loss (black). **(H)** Target signals, E estimates and I estimates in three input dimensions (three top rows), spike trains (fourth row) and the instantaneous estimate of the firing rate of E and I populations (bottom) in a simulation trial, with significant jitter of recurrent connectivity (jittering strength of 0.5, see Methods). In spite of a relatively strong jittering, the network shows excellent encoding of the target signal. All statistical results were computed in 100 simulation trials of duration of 1 second. Other parameters are in Table 1. This figure is related to Fig. 4 in the main paper.

**Figure S4.**
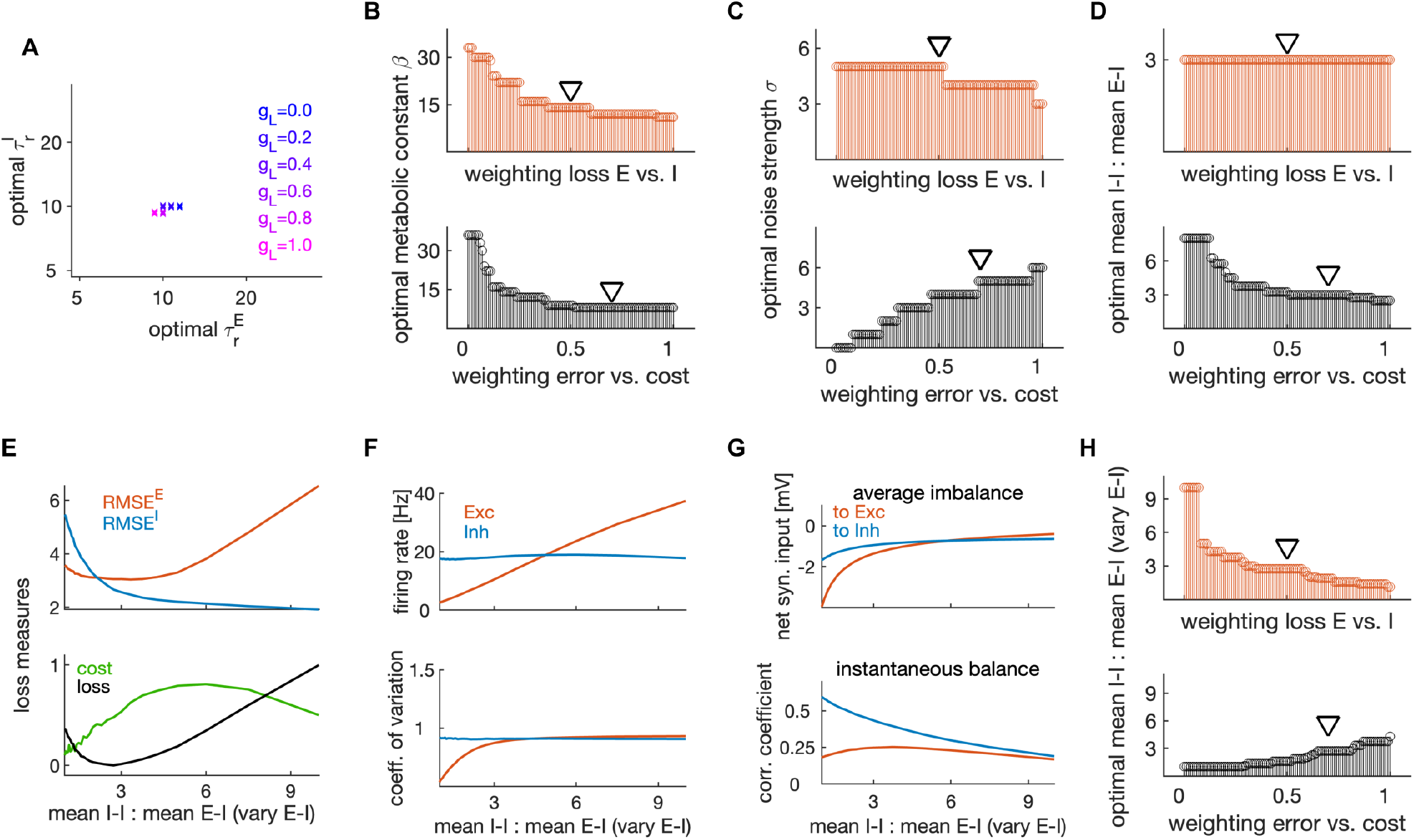
Dependence of optimal parameters on weighting of the encoding error and the metabolic cost and analysis of mean ratio of I-I to E-I connectivity by varying the number of E neurons. **(A)** Optimal set of time constants of E and I neurons 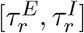 for different weightings between the error and the cost when computing the loss. Optimal time constants show little dependency on this weighting. **(B)** Top: Optimal metabolic constant as a function of the weighting of the average loss of E and I cell type. Bottom: Same as on top, as a function of the weighting between the error and the cost. Black triangles mark weightings that are typically used to estimate optimal model efficiency. **(C)** Same as in **B**, as a function of noise strength. **(D)** Same as in **B**, as a function of the optimal ratio of I-I to E-I connectivity. This analysis was performed by varying the number of I neurons while the number of E neurons stays fixed. **(E)** Top: Encoding error (RMSE) of the E (red) and I (blue) estimates as a function of mean I-I to E-I connectivity. The ratio was varied by changing the number of E neurons and keeping the number of I neurons fixed at a value specified in Table 1. Bottom: Same as on top, showing the normalized cost and average loss. **(F)** Same as in **E**, showing, the average firing rate (top), and average coefficient of variation (bottom) in E and I cell type. **(G)** Same as in **E**, showing the average imbalance and instantaneous balance of synaptic currents in E and I neurons. **(H)** Same as in **D**, for the optimal ratio measured by varying the number of E neurons. All results were computed in 100 trials of duration of 1 second for each trial. For other parameters, see Table 1. This figure is related to Fig. 7 in the main paper.

**Figure S5.**
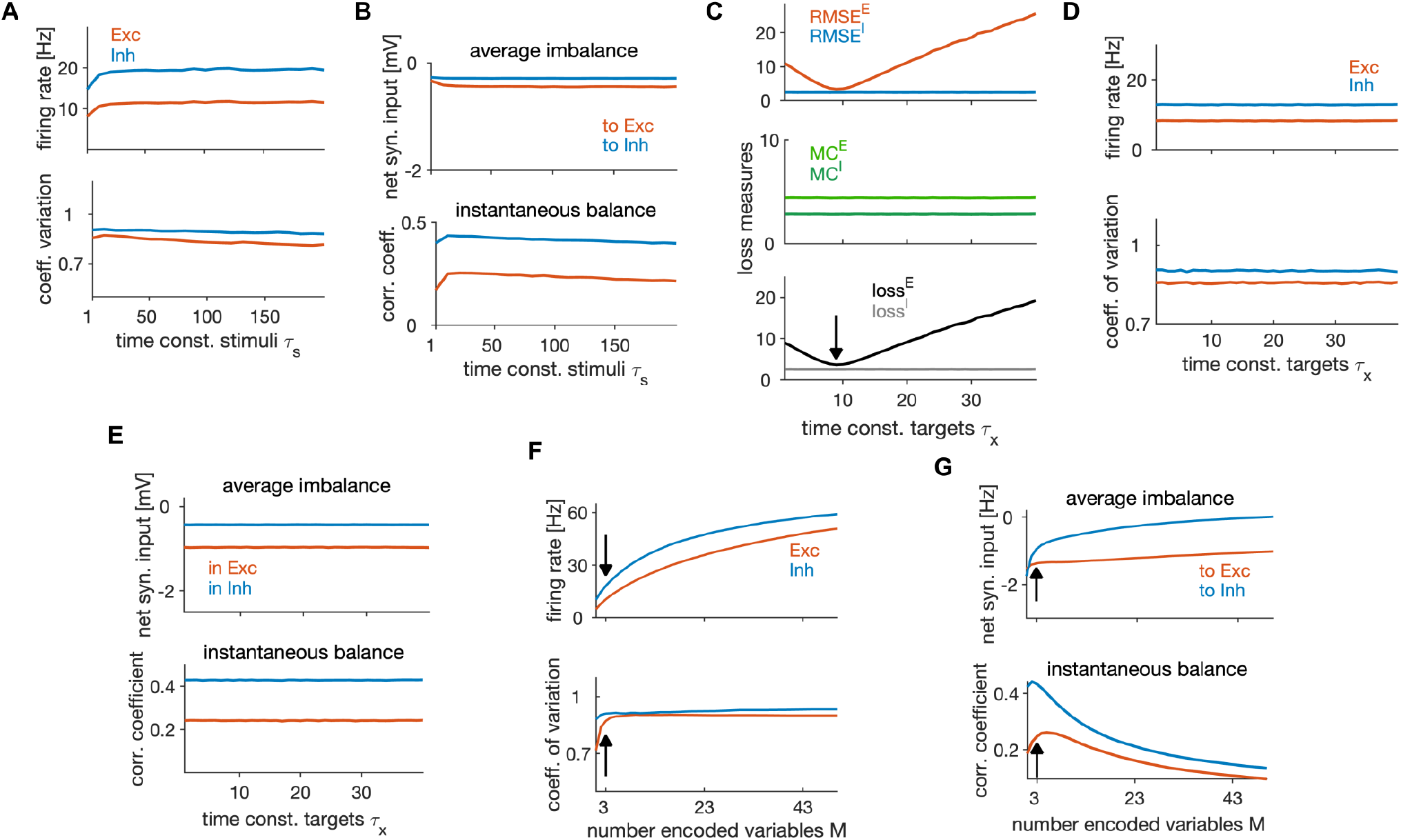
Effect of stimulus properties on efficient neural coding and dynamics. **(A)** Average firing rate (top), and average coefficient of variation (bottom) in E and I cell type, as a function of the time constant of the stimulus features *τ*_*s*_. All stimulus features have the same time constant. **(B)** Average imbalance (top) and instantaneous balance (bottom) as a function of the time constant of stimuli *τ*_*s*_. **(C)** Top: RMSE of E (red) and I (blue) estimates as a function of the time constant of the targets *τ*_*x*_. All targets have the same time constant. Middle: Metabolic cost in the E and I population. Bottom: Average loss in the E and I population. Black arrow indicates the minimum loss and therefore the optimal time constant. **(D-E)** Same as in **A-B**, as a function of the time constant of the targets *τ*_*x*_. **(F-G)** Same as in **A-B**, as a function of the number of encoded variables *M*. All results were computed in 100 trials of duration of 1 second. For parameters, see Table 1. This figure is related to Fig. 8 in the main paper.

